# Lipid Nanoparticles from *L. meyenii* Walp Mitigate Sepsis through Multimodal Protein Corona Formation

**DOI:** 10.1101/2024.11.13.623417

**Authors:** Junsik J. Sung, Jacob R. Shaw, Josie D. Rezende, Shruti Dharmaraj, Andrea L. Cottingham, Mehari M. Weldemariam, Jace W. Jones, Maureen A. Kane, Ryan M. Pearson

## Abstract

**Background:** Plant-derived nanoparticles (PDNP) are nano-sized particles isolated from various edible plants that contain bioactive components involved in regulating cellular immune responses against pathogenic intrusion and inflammation.

**Purpose:** This study describes a novel PDNP derived from *Lepidium meyenii* Walp (maca) that efficiently captures pro-inflammatory cytokines and acute phase proteins in its protein corona to enhance survival in two representative lethal models of sepsis.

**Methods:** Lipid nanoparticles were isolated from maca (MDNP) and triacylglycerols and phytoceramides were identified as major constituents using lipidomics. The physicochemical properties of MDNPs were determined, anti-inflammatory effects of MDNP were evaluated using *in vitro* models and *in vivo* using endotoxemia and cecal ligation and puncture (CLP) polymicrobial sepsis models. Proteomic analysis of MDNP in healthy or LPS-induced inflammatory plasma was used to determine the composition and inflammatory pathways modulated due to the MDNP protein corona.

**Results:** *In vitro* studies showed that MDNP were non-toxic, reduced macrophage activation, and effectively sequestered pro-inflammatory cytokines to mitigate NF-*κ*B activity under lipopolysaccharide (LPS) stimulation. In a pre-established LPS-induced endotoxemia model, MDNP-treated mice showed significantly reduced systemic pro-inflammatory cytokines and enhanced survival. Untargeted proteomics and pathway analysis of the MDNP protein corona identified an enrichment in acute phase proteins in MDNP-LPS plasma coronas. MDNP treatment also significantly improved survival in the CLP sepsis model in the absence of antibiotics.

**Conclusion:** This work identified MDNP as an efficient, plant-derived lipid NP that broadly sequesters and neutralizes a compilation of inflammatory mediators in their coronas, offering multimodal therapeutic potential for treating inflammatory diseases.

## Introduction

Edible plant-derived nanoparticles (PDNP) are nanostructured and membrane-enveloped vesicles secreted by plant cells that serve as carriers of various endogenous bioactive substances. Although previously perceived as cellular debris, PDNP are recognized as crucial entities that regulate cell-cell communication and immune responses against pathogens (Shao et al., 2018). The evolving understanding of the biogenesis of PDNP has been described in previous literature (Chafe, 1969; Robarbs and Kidwai, 1969). Briefly, the process initiates with the formation of a trans-Golgi network or early endosome. Plant cells then generate the matured multivesicular endosome that integrate and fuse with the plasmalemma, releasing PDNP through the inward budding of multivesicular endosomes (Li et al., 2018). There are several studies pointing to the potential therapeutic benefits of PDNP, which in some cases demonstrate anti-inflammatory (Yang et al., 2021), antioxidant (Perut et al., 2021), and anti-cancer activity (Wang et al., 2013). This emerging class of naturally derived nanovesicles offers highly advantageous features for use as nanomedicines, exhibiting negligible toxicity or immunogenicity, efficient cellular uptake, and having capacity to deliver a variety of therapeutic agents through re-engineering (Fang et al., 2024).

Cytokine secretions and the acute phase response (APR) are early, systemic immune reactions that develop within minutes to hours as part of the body’s early defense mechanism to injury, infection, or immune challenge (Chen et al., 2018; Markanday, 2015a). The APR is triggered by the release of pro-inflammatory cytokines such as interleukin-6 (IL-6), tumor necrosis factor-alpha (TNF-α), interleukin-1b (IL-1β), and others by macrophages, neutrophils, and various immune cells at the site of inflammation (Tanaka et al., 2014). This response triggers the production of acute phase proteins (APP), which are typically undetectable under healthy conditions but increased 10-to 1000-fold during inflammation (Ansar and Ghosh, 2016). These proteins play roles in modulating inflammation, enhancing pathogen clearance, and promoting tissue repair (Ehlting et al., 2021). The APR is generally recognized as beneficial for restoring homeostasis disturbed by such injuries, however, APP have been described to elicit a range of functions to induce pro- or anti-inflammatory responses. During severe inflammatory responses like sepsis, a dysregulated APR can potentiate the inflammatory response leading to coagulopathy, organ failure, and even death (Castellheim et al., 2009; Markanday, 2015b; Zhang and Ning, 2021).

Several groups have developed strategies to mitigate the inflammatory response by sequestering pro-inflammatory cytokines (Lasola et al., 2020). Multiple studies highlight that the use of membrane-coated nanoparticles that mimic biological cell membranes to trap and neutralize these inflammatory cytokines (Khatoon et al., 2022). By acting as macrophage decoys, these nanoparticles can be developed to bind and neutralize endotoxins, pro-inflammatory cytokines, and even venoms that would contribute to negatively contribute to survival outcomes (Thamphiwatana et al., 2017).

Similarly, neutrophil membrane-coated nanoparticles are produced using neutrophils isolated from mouse bone marrow after LPS stimulation (Zhang et al., 2024). The process involves membrane extraction, incubation, and extrusion or sonication with polymeric NP cores. These membrane-camouflaged nanoparticles reduce inflammation by binding and neutralizing endotoxins and cytokines such as IL-6, and TNF-α (Shen et al., 2019; Thamphiwatana et al., 2017). A study showed that IL-6 can be effectively captured and neutralized using chitosan/hyaluronic acid polymeric nanoparticle that have antibodies immobilized on the surface (Lima et al., 2018). Moreover, telodendrimers were developed to bind LPS and pro-inflammatory cytokines through controlling hydrophobic and charge-based interactions (Shi et al., 2020a), and poly(ethylene glycol) (PEG) hydrogels were developed to capture and neutralize histones to improve sepsis survival (Koide et al., 2021a). Despite these advancements, the complex synthesis and formulation required for these designs to sequester pro-inflammatory mediators presents a significant challenge to clinical translation. As a result, there is critical need to develop simpler anti-inflammatory strategies that can sequester multiple inflammatory mediators to mitigate pro-inflammatory responses to augment sepsis survival.

*Lepidium meyenii* Walp, also referred to as maca, is a biennial root plant indigenous to the Peruvian Andes. Maca possesses high contents of fiber, amino acids, fatty acids, and other essential nutrients, while also containing various bioactive compounds that are involved in integral cellular activities within plants (Wang and Zhu, 2019). Several studies have shown that maca alleviates fatigue (Zhu et al., 2022), oxidative stress (Fei et al., 2022), tumor formation (Cao et al., 2023), and inflammation (Yang et al., 2023). However, studies have focused on the raw material or crude extract, which contains mixture of different active ingredients, to evaluate in disease models (Fei et al., 2020; Tenci et al., 2017). Despite these known advantages, maca remains relatively underexplored.

This investigation demonstrates the isolation, characterization, and therapeutic development of a PDNP isolated from maca root, termed maca-derived lipid nanoparticle (MDNP), for the treatment of severe inflammation as exemplified using *in vitro* and *in vivo* models of lipopolysaccharide (LPS)-induced endotoxemia and polymicrobial sepsis. MDNP were isolated and characterized for physicochemical properties, as well as lipid composition using lipidomic analysis. The toxicity profile of MDNP was then established prior to determining their uptake profile and anti-inflammatory properties. *In vitro*, MDNP efficiently sequestered multiple pro-inflammatory cytokines, which led to comprehensive *in vivo* assessment of their biodistribution and therapeutic activity. Therapeutic administration of MDNP to LPS-challenged mice led to significant reductions in plasma pro-inflammatory cytokines, reductions in inflammation-induced organ damage, and improved survival. When nanoparticles are introduced into a biological fluid, they are rapidly covered by a layer of biomolecules known as the biomolecule corona (or protein corona) (Pearson et al., 2014; Shaw and Pearson, 2022). To identify the mechanism by which MDNP elicited its anti-inflammatory effects, we performed untargeted proteomics analysis of the MDNP protein corona to uncover the inflammatory mediators sequestered by MDNP and corresponding pathways and upstream regulators modulated. MDNP was found to sequester and neutralize a variety of APP, which promote the propagation of pro-inflammatory immune responses, in addition to pro-inflammatory cytokines. Lastly, the efficacy of MDNP was assessed using a clinically relevant mouse model of polymicrobial sepsis and found to significantly increase survival. These results demonstrate the potential of MDNP as an abundant, cost effective, naturally derived therapeutic agent for use as a multimodal intervention for a variety of inflammatory diseases due to their ability to sequester a broad spectrum of inflammatory mediators.

## Results

### Isolation and characterization of maca-derived lipid nanoparticles (MDNP)

PDNPs were isolated from maca juice using differential ultracentrifugation and density gradient sucrose gradient centrifugation (**Figure 1**) (Sung et al., 2019). MDNP accumulated at the interface of 20/35% sucrose gradient, and the recovery was approximately 4 mg per 30 g of maca powder (**Figure 1B,C**). TEM revealed that MDNP displays a spherical morphology, and their core structure was consistent with that of solid lipid nanoparticles (**Figure 1D**). Size and zeta potential of MDNP measured an average of 174.5 nm and −9.6 mV, respectively (**Figure 1E**). We next assessed the stability of MDNP at multiple temperatures. MDNP could tolerate a freeze and thaw cycle and were stable at 4°C for up to 6 days, the maximum length of time tested. All different temperature storage conditions kept MDNP stable except for room temperature (RT), where after 3 days the size was significantly increased (**Figure 1F**). The sizes and zeta potential of MDNP were also stable upon lyophilization (**Supplementary Figure S1**). Differential scanning calorimetry (DSC) analysis determined the summit of melting peak of MDNP, which was measured as 74.3°C (**Figure 1G**).

**Figure 1.**
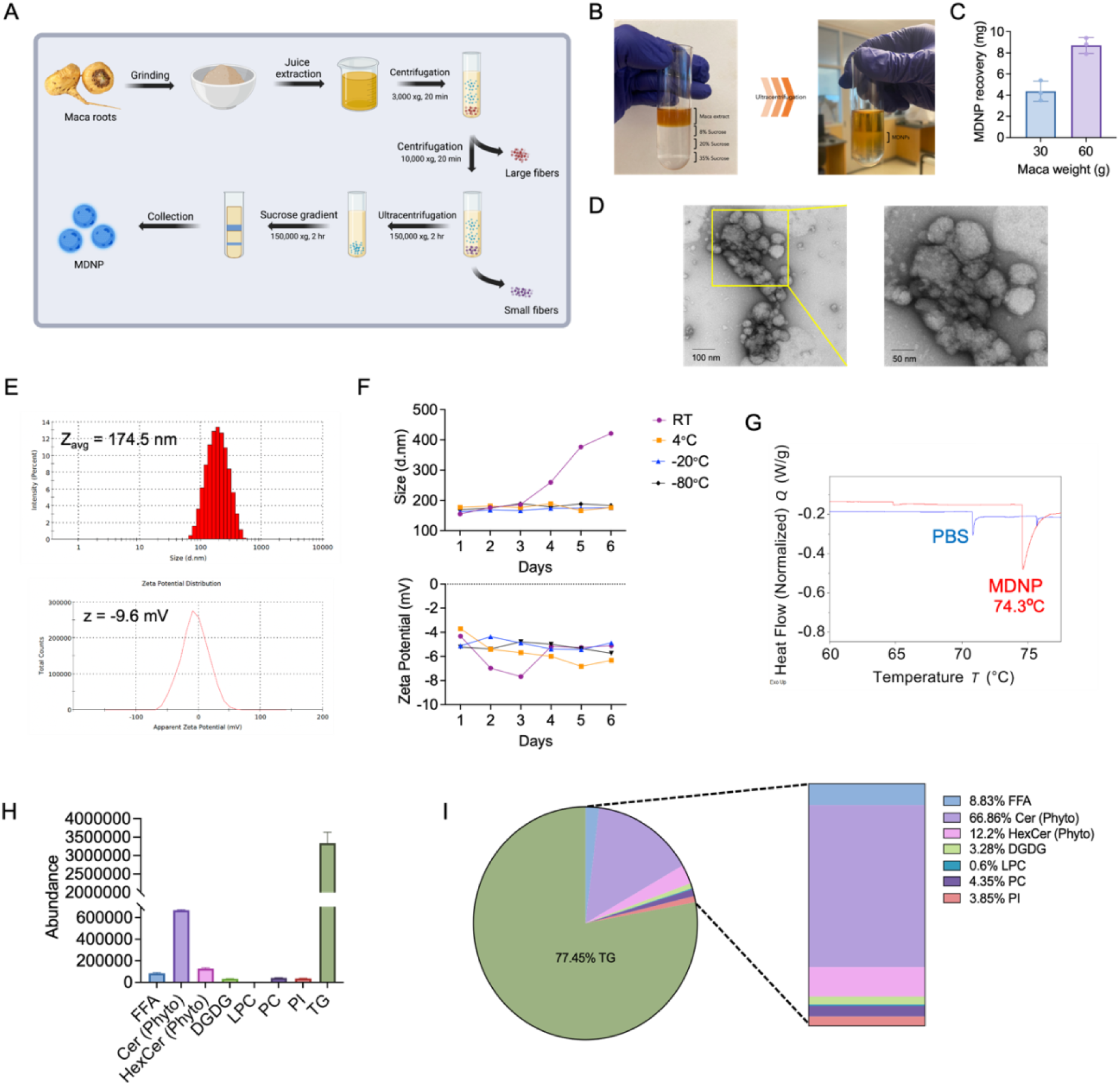
Isolation and characterization of MDNP. (A) Schematic representation of the isolation process for maca-derived lipid nanoparticles (MDNP). (B) Before and after images of MDNP isolation. MDNP were isolated and purified from maca extract by sucrose density gradient purification. (C) MDNP recovery from pure maca powder. (D) A representative TEM image of MDNP. (E) Room temperature measurement of size (174.5 nm) and zeta potential (−9.6 mV) of freshly isolated MDNP. (F) Storage stability test of MDNP under various storage conditions up to day 6. (G) Heat flow curve of MDNP measured by differential scanning calorimetry showing the summit of melting peak at 74.3 °C. (H) Lipid extraction and component analysis by liquid chromatography coupled to high resolution LC-MS/MS revealed eight different lipids (TG, Cer, HexCer, DGDG, LPC, PC, PI and FFA). Bar graph shows abundance of lipids. (I) Pie graph shows the whole ratio of each component and the vertical slice magnifies the smaller portion of composition excluding TG. N=3/group.

To determine the lipid composition of MDNP, untargeted high throughput lipidomics was performed. A bar graph (**Figure 1H**) and a pie/vertical slice chart (**Figure 1I**) shows the total lipid abundance and ratio in percentage of each component in MDNP. The lipidomics identified a total of eight individual lipids in four categories including sphingolipids, galactolipids, neutral lipids, and phospholipids (**Supplementary Figure S2**). The most prevalent species within the MDNP was the triglyceride (TG) subclass with 51 identifications (77.45%), followed by phytoceramide (Cer (Phyto); 14.77%), phytohexosylceramide (HexCer (Phyto)) (2.7%), free fatty acids (FFAs) (1.95%), digalactosyldiacylglycerol (DGDG) (0.72%), phosphatidylcholine (PC) (0.96%), phosphatidylinositol (PI) (0.85%), and lysophosphatidylcholine (LPC) (0.13%).

The complete protein compositional analysis of MDNP using UPLC-MS/MS revealed undetectable levels of proteins that could be linked to plant databases from Uniprot and the Plant Proteome Database. This aligns with our lipidomics results indicating that MDNP is primarily composed of lipids, with minimal or no associated protein content. Overall, these data demonstrate the successful isolation, purification, and characterization of MDNP from bulk maca extract.

### *In vitro* cytotoxicity, internalization, and therapeutic effect of MDNP

To analyze potential toxicity, BMDM were treated with MDNP (1 mg/mL) for 8 hours prior to flow cytometry analysis to measure viability and apoptosis (**Figure 2A**). MDNP treatment did not affect the cell viability or induce apoptosis. Furthermore, we observed a time-dependent cellular uptake of Cy5.5-labeled MDNP using BMDM (**Figure 2B**). The quantitative measure of mean fluorescence intensity (MFI) from the confocal microscopy images can be found in **Supplementary Figure S3**. These findings demonstrate the MDNP are non-toxic and efficiently internalized by BMDM.

**Figure 2.**
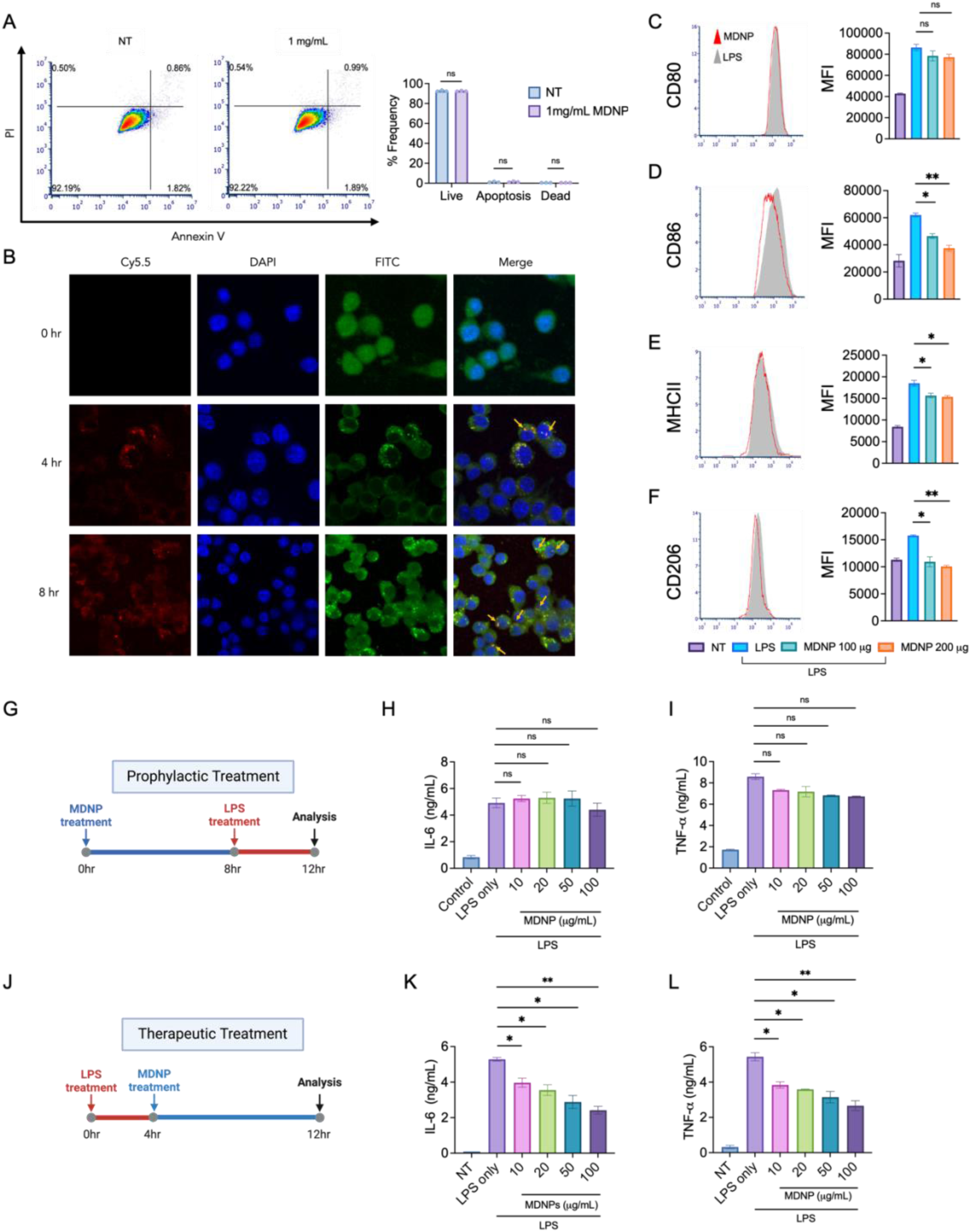
*In vitro* cytotoxicity, cellular internalization, immunomodulation and therapeutic effects against LPS-induced inflammation. (A) PI/Annexin V assay by flow cytometry displayed that treatment with higher concentration (1 mg/mL) of MDNP did not induce any significant toxicity in BMDM. Quantitative analysis of live, apoptotic, and dead cells after MDNP treatment is accompanied. (B) Cy5.5 labeled MDNP were gradually internalized by BMDM after 4 and 8 hrs. Cells were stained with DAPI and FITC-PI to visualize the nucleus and F-actin, respectively. (C-F) Immunomodulatory activity of MDNP in LPS inflamed BMDM showed reduction in M1 (CD80, CD86, MHCII) and M2 (CD206) markers (G) A timeline for prophylactic treatment. Prophylactic approach did not make significant change in levels of (H) IL-6 and (I) TNF-⍺. (J) A timeline for therapeutic treatment. Therapeutic treatment displayed significant reduction of (K) IL-6 and (L) TNF-⍺. Data are expressed as mean ± SD (n=5). Data are expressed as mean ± SD (n=3). *P<0.05, **p<0.05 versus LPS control group.

BMDM were treated with LPS followed by MDNP to assess their ability to mitigate immune activation. The observed decrease in CD80 (reduced but not significant), CD86, and MHCII expression following MDNP treatment suggests that MDNP reduced macrophage activation, contributing to a decreased pro-inflammatory response (**Figure 2C-E**). Modulation of macrophage polarization is one of the strategies to modulate inflammation, support tissue regeneration and reestablish disrupted homeostasis(Parisi et al., 2018). We also measured a reduction in CD206 as a marker of anti-inflammatory M2-like macrophages, suggesting MDNP induced a shift in macrophage phenotype towards a more undifferentiated or M0-like state (**Figure 2F**). These results indicate that MDNP can impact the activation status of macrophages suggesting that they aid in shifting their profile towards a more balanced and neutral immune response.

To assess the anti-inflammatory effects of MDNP, we investigated its prophylactic and therapeutic activity by treating MDNP on LPS-challenged BMDM. In the prophylactic setting, MDNP was treated for 8 hr prior to 4 hr LPS exposure (**Figure 2G**). However, it did not significantly attenuate the release of pro-inflammatory cytokines (IL-6 and TNF-α) (**Figure 2H,I**). In the therapeutic setting, BMDM were challenged with LPS for 4 hrs prior to the treatment of MDNP for 8 hr (**Figure 2J**). The result shows that pro-inflammatory cytokines including IL-6, and TNF-α induced by LPS were significantly reduced, whereas IL-1β, and IFN-γ were only slightly reduced (**Supplementary Figure S4**). Furthermore, additional testing was conducted on the crude extract after MDNP isolation to determine if it possesses any anti-inflammatory efficacy; however, no cytokine reduction was detected (**Supplementary Figure S5**). These results suggested that MDNP can reduce pro-inflammatory responses, and the anti-inflammatory component of maca is the purified MDNP.

### Sequestration of cytokines and mitigation of NF-κB activation

Interleukin-6 (IL-6) and tumor necrosis factor-α (TNF-α) are well known to be involved in pathogenesis of chronic inflammation, autoimmune diseases, and cancer (Hirano, 2021; Popa et al., 2007). Utilizing data from the *in vitro* anti-inflammatory assay, an investigation was conducted to confirm whether MDNP possess the capability to sequester pro-inflammatory cytokines in the absence of cells. In a time-dependent experiment, BMDM were treated with 100 ng/mL LPS for 4 hrs, then the supernatant rich in inflammatory cytokines was incubated with 100 µg/mL MDNP for up to 24 hrs (**Figure 3A**). ELISA analysis of these supernatants indicated a gradual decrease in IL-6 levels, with a maximum of 48% reduction after 8 hrs (**Figure 3B**). A similar trend was observed for TNF-α levels, with a 28% reduction over the same time period (**Figure 3C**). Optimization of MDNP concentration revealed concentration-dependent cytokine sequestration properties of MDNP and 100 µg/mL MDNP exhibited the most significant sequestration of IL-6 and TNF-α, whereas PLGA nanoparticle (as control) did not display any cytokine sequestration properties (**Supplementary Figure S6**). We also confirmed the ability of MDNP to sequester recombinant IL-6 to exclude the possibility of other proteins contributing to cytokine removal *in vitro* (**Figure 3D,E**). These findings confirm that MDNP can sequester pro-inflammatory cytokines to mitigate inflammatory responses.

**Figure 3.**
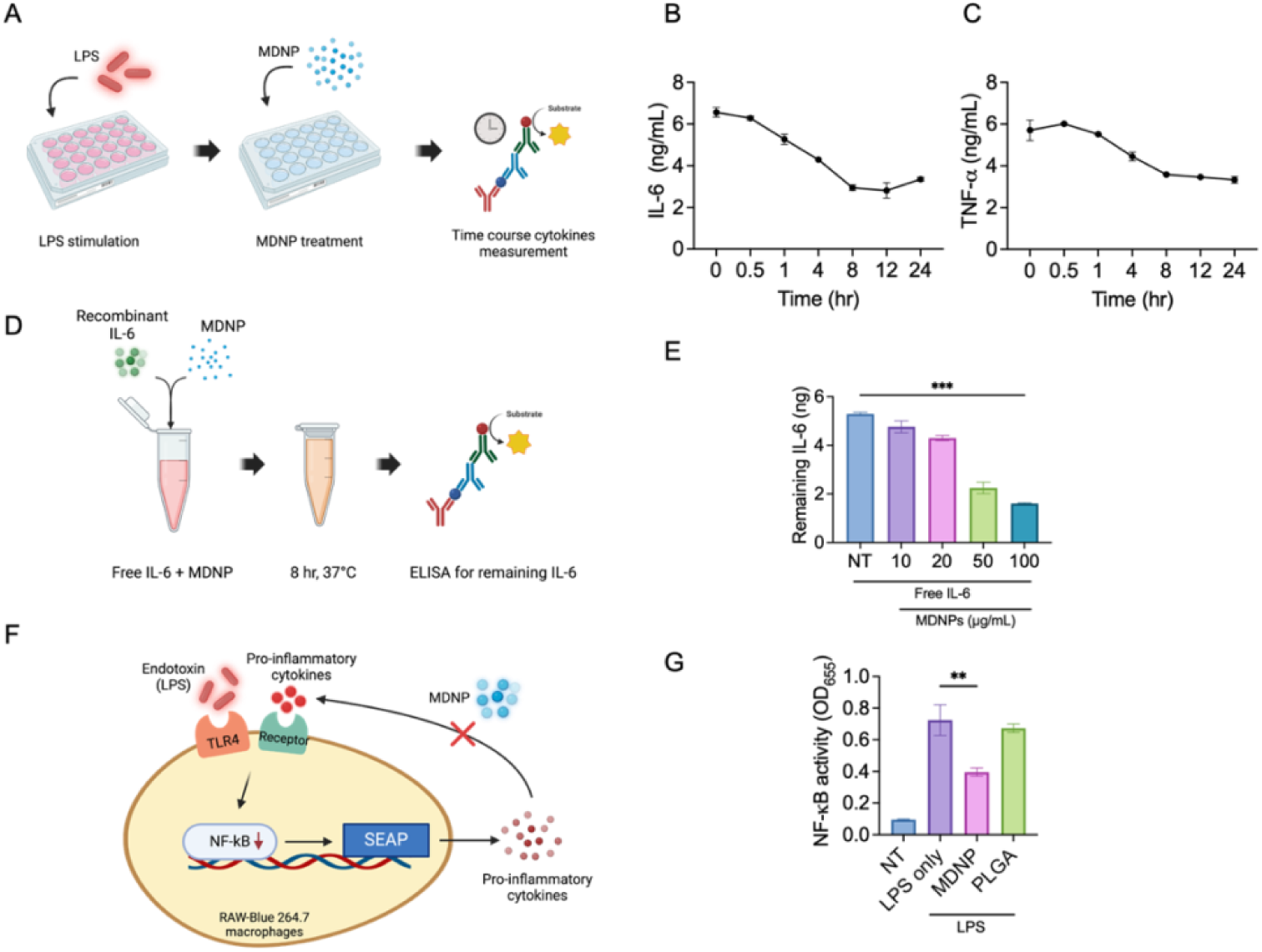
*In vitro* sequestration of pro-inflammatory cytokines from LPS-induced bone marrow derived macrophages. (A) Time course reduction of LPS-stimulated pro-inflammatory cytokine by treatment of MDNP. 8 hr time point exhibited the most significant reduction for both (B) IL-6 and (C) TNF-α. (D) Recombinant IL-6 was sequestered through incubation with MDNP ex vivo without cellular interaction. (E) 100 µg/mL MDNP was the most effective concentration to remove IL-6 protein. (F) Expression of NF-*κ*B in in RAW-Blue 264.7 cell is suppressed by the treatment of MDNP. (G) NF-*κ*B activity level is downregulated after treatment of MDNP. PLGA was used as a control to show different effect on NF-*κ*B. All data are expressed as means ± SD (n=5). **p<0.05, ***p<0.005, and ****p<0.0005 versus LPS or NT control group.

The innate immune responses that are triggered by LPS are mediated by Toll-like receptor (TLR)-4 and subsequent activation of the transcription factors nuclear factor kappa B (NF-*κ*B) and activator protein 1 (AP-1) (Nyati et al., 2017). Upon encountering various PAMPs and DAMPs, macrophages undergo rapid activation and secrete a diverse range of cytokines and chemokines including IL-6 and TNF-α (Liu et al., 2017). In this experiment, RAW-Blue cells, a NF-*κ*B macrophage reporter cell line that releases secreted alkaline phosphatase (SEAP) after activation, were used to gauge MDNP-mediated effects following LPS stimulation (**Figure 3F**). The expression of IL-6 is tightly regulated at multiple levels, including gene transcription, mRNA translation and mRNA degradation levels. Furthermore, NF-*κ*B activation was examined by exposing LPS-activated RAW-Blue cells to 100 µg/mL MDNP for 1 hr (**Figure 3G**). PLGA was used as a control and were unable to reduce NF-*κ*B activation. The results demonstrate a significant decrease in NF-*κ*B activity, suggesting that MDNP possess the capacity to mitigate pro-inflammatory cell signaling and reduce macrophage activation.

### *In vivo* biodistribution and anti-inflammatory effects of MDNP

For the *in vivo* biodistribution study, MDNP were labeled with Cy5.5 and administered intraperitoneally (IP) or intravenously (IV) into LPS-challenged mice (5 mg/kg LPS dose) twice at an MDNP dose of 2 mg/injection at 0.5 and 2 hr (**Figure 4A**). At 4 hr, mice were euthanized to image the biodistribution of Cy5.5-MDNP in different organs including liver, spleen, heart, kidney, and lung with an *in vivo* imaging system (IVIS). Most of the Cy5.5-labeled MDNP were delivered to the liver and kidney by both IP and IV injection, and very low amounts were detected in the spleen, heart, and lung (**Figure 4B, C**).

**Figure 4.**
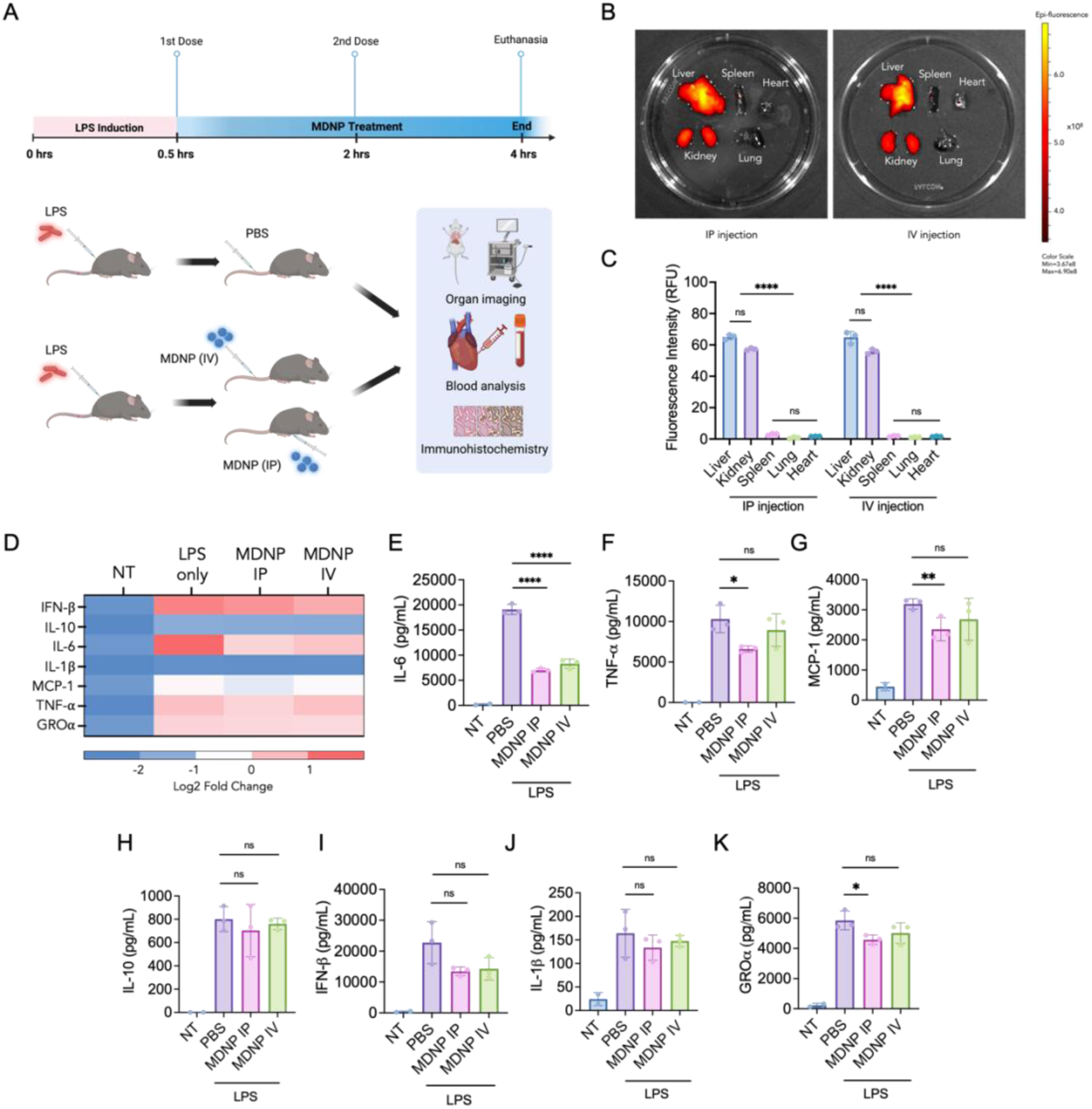
Route of administration biodistribution and anti-inflammatory effects of MDNP (A) Timeline of MDNP treatments following LPS challenge. The schematic illustrates the IP and IV injection performed on randomly selected C57BL/6J mice groups (LPS only, LPS+IP injection, and LPS+IV injection, n=3 per group). (B) Biodistribution of fluorescently labeled MDNP in mouse organs (liver, kidney, spleen, lung, and heart). (C) Fluorescence intensity measurement of MDNP biodistribution. (D) Heatmap of pro-inflammatory cytokine expression including: (E) IL-6, (F) TNF-α, (G) MCP-1, (H) IL-10, (I) IL-1β, (J) IFN-β, (K) GROα from mice plasma samples. Data are expressed as mean ± SD (n=3). *P<0.05, **p<0.05, ***p<0.005, and ****p<0.0005 versus PBS control group.

We also investigated how the route of administration (IP versus IV) affected the *in vivo* therapeutic effects in mice. Mice were challenged with 5 mg/kg LPS IP followed by treatment with 2 mg MDNP at 0.5 hr and 2 hr post-LPS treatment. After 4 hrs, mice were euthanized to collect organs samples and plasma (**Figure 4A**). Plasma samples were processed using a MAGPIX Luminex multiplexing immunoassay system. This system was employed to analyze the expression level of a 7-plex panel of pro-inflammatory cytokines which included IL-6, TNF-α, MCP-1, IL-10, IL-1β, IFN-β, and GROα. The heatmap demonstrates overall reduction of pro-inflammatory cytokines for both routes (**Figure 4D**). The analysis revealed a significant reduction in IL-6 level following both administration routes (**Figure 4E**), and a notable reduction in TNF-α, MCP-1, and GROα for IP administered MDNP (**Figure 4F-K**). IV administration of MDNP also reduced systemic cytokine levels, however it was slightly less effective. We further evaluated blood biochemistry of albumin, alanine transaminase, creatine, aspartate transferase, globulin, total protein, and blood urea nitrogen. Although minor reductions were observed for MDNP-treated mice, there was no significant alteration in levels of blood biochemistry parameters compared to LPS-treated controls (**Supplementary Figure S7**). The non-significant reductions in blood biochemistry following MDNP-treatment could be due to the pre-established severe inflammatory response, that could not be completely reversed by the endpoint of the study.

### Treatment of MDNP accelerated organ recovery and improved survival

Histological analysis was used to evaluate the level of inflammation in tissues isolated from experiments conducted in **Figure 4**. H&E staining of five different organs including liver, kidney, spleen, lung, and heart are shown in **Figure 5**. LPS-treated liver showed severe immune cell infiltration, tubular necrosis was detected in the kidney, focal necrosis, and dysregulation in white pulp of spleen was observed, and extensive clots in the alveoli of lung appeared, but no significance of heart injury was measured. On the contrary, the IV and IP administered MDNP groups displayed notable improvements in histology scores when compared to the LPS group (**Figure 5B-F**). Based on the overall safety profile and reductions in pro-inflammatory cytokines, MDNP were used to further evaluate survival *in vivo*. Mice were challenged with lethal dose of 20 mg/kg LPS in sterile PBS intraperitoneally. Then, mice were monitored after two doses of 2 mg MDNP (**Figure 5G**). As a result, 60% of MDNP treated mice survived, whereas 0% of the LPS treated (control) mice survived (**Figure 5H**).

**Figure 5.**
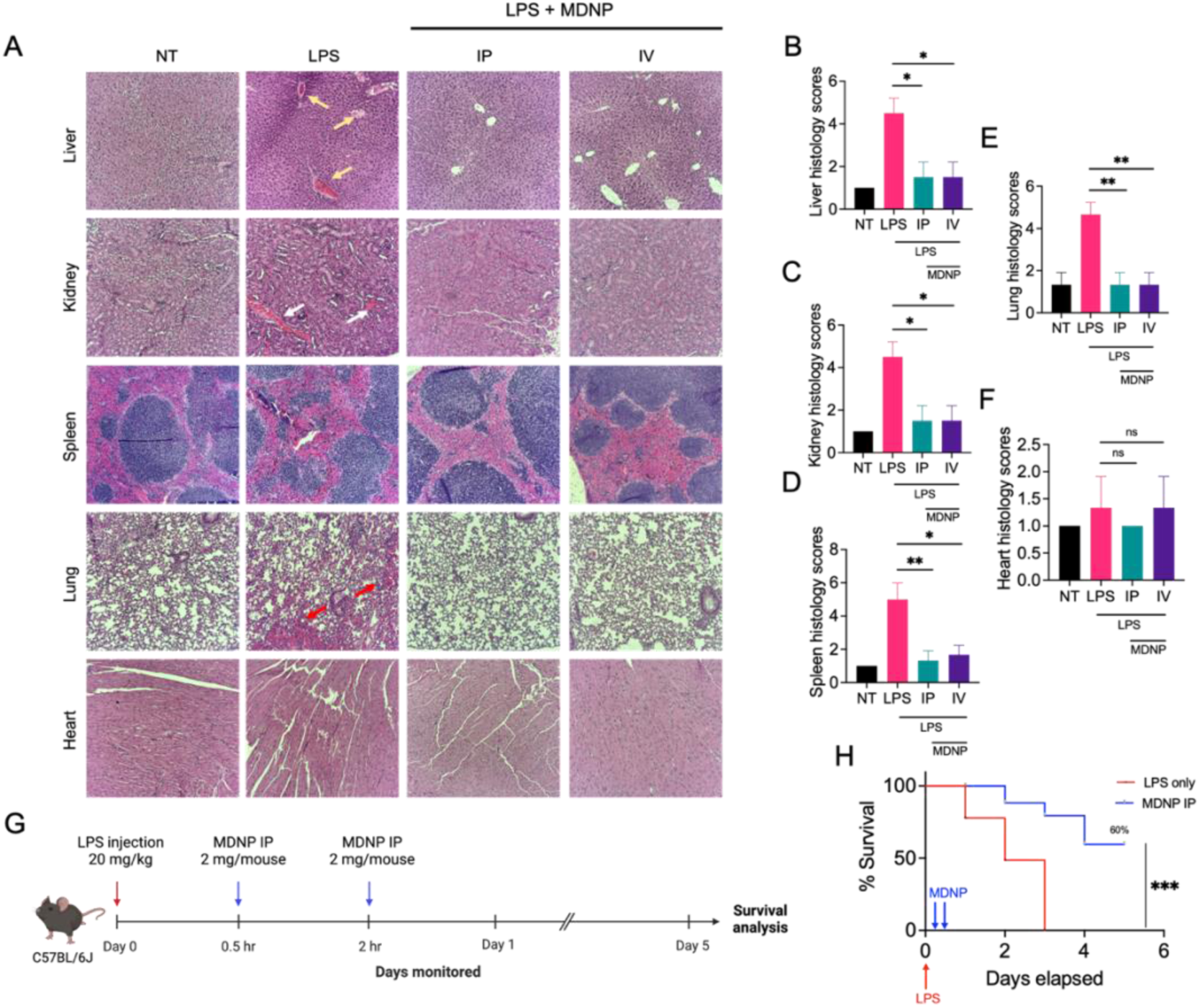
Representative images of organ recovery and survival effect by treatment of MDNP during LPS-induced sepsis. (A) H&E staining of liver, kidney, spleen, lung, and heart. The LPS treated group images shows presence of neutrophil infiltration (yellow arrow) in liver, tubular necrosis (white arrow) in the tubules in kidney, focal necrosis and dysregulation in white pulp of spleen, appearance of extensive clots (red arrow) in the alveoli in lungs, while no noticeable damage in heart. On the contrary, mice organs administered with IP and IV displayed notably reduced level of organ damage of organ damage. (B-F) Quantification of H&E staining. Levels of injury scores were calculated. Data are expressed as mean ± SD (n=3). *P<0.05, **p<0.05 versus LPS control group. (G) A schematic of survival study of LPS-treated and LPS+MDNP-treated C57BL/6J mice (n=10/group). (H) The survival analysis graph displays a significant difference between LPS only and the MDNP treatment group. Kaplan-Meier curves and a log-rank test was performed to compare the survival rates. ***p<0.0005.

### Proteomic profiling of MDNP protein corona

The ability of MDNP to reduce systemic cytokines and promote LPS mouse survival prompted an exploration of the composition and types of proteins sequestered by the MDNP. Adsorbed proteins were characterized by forming a protein corona on MDNP *ex vivo* by incubating with either healthy or LPS-treated mouse plasma (MDNP-H or MDNP-LPS, respectively) (**Figure 6A**). Proteomics analysis of these coronas identified 297 total proteins (**Supplementary Table S1**), with 270 common between the two conditions, 1 distinctive for healthy, and 26 for LPS-treated (**Figure 6B**). **Table 1** presents a list of uniquely adsorbed proteins from each group. Among these, CD14, LCN2 (lipocalin-2), APCS (Serum Amyloid P component), FABP4 (Fatty Acid Binding Protein 4), NGP (Neutrophilic Granule Protein), and HSP90AA (Heat Shock Protein 90 alpha) are known to play roles in inflammation and categorized as APP excluding NGP. The heatmap in Figure 6C shows the differential expression of total 297 detected proteins. Comparing MDNP-H to MDNP-LPS, highlighted significant changes in protein abundance, with 49 proteins being significantly upregulated and 100 proteins downregulated (**Figure 6D**). Haptoglobin (Hp), which is known for its pro-inflammatory properties(Shen et al., 2011), was found to be the most differentially abundant protein in the MDNP-LPS corona. The top 15 proteins (over 20-fold change) identified in the MDNP-LPS group are shown in **Supplementary Figure S8**. Among these, Hp, Seprina3n, and Saa1 are well-known for their pro-inflammatory roles in regulating inflammatory process and promoting the release of inflammation cytokines. Their levels can increase dramatically during inflammation(Massart et al., 2020). Figure 6E lists the canonical pathways of MDNP-LPS coronas from ingenuity pathway analysis (IPA). APR signaling was most highly associated with MDNP-LPS corona. The network diagram in Figure 6F illustrates the complex but close interaction of APP and these upstream pro-inflammatory regulators within the APR signaling pathway. The highlighted connections between APR shows its direct interaction with the APP sequestered on MDNP-LPS coronas (Hp, Saa1, SerpinA3N) and a subset of upstream regulators (IL-6, TNF, IL1B, AGT, HNF1A). Importantly, IL-6, TNF and IL1B play pivotal roles in either indirectly or directly activating these APP. Figure 6G illustrates the result of *ex vivo* cytokine sequestration assays using MDNP. It demonstrates that MDNP retains the ability to bind and remove pro-inflammatory cytokines, such as IL-6 and TNF-α, from the LPS plasma. It is also important to point out that protein corona formed on the surface of MDNP did not activate any pro-inflammatory responses when treated on BMDM, supporting that the formation of the corona effectively neutralizes the pro-inflammatory responses of these various mediators (**Supplementary Figure S9**). Figure 6H summarizes the proposed mechanism of MDNP by forming a multimodal protein corona comprised of pro-inflammatory cytokines and APP and neutralization. This inhibits the key inflammatory pathways, blocking the production of pro-inflammatory upstream regulators, ultimately reducing pro-inflammatory cytokines, improving organ function, and increasing survival.

**Figure 6.**
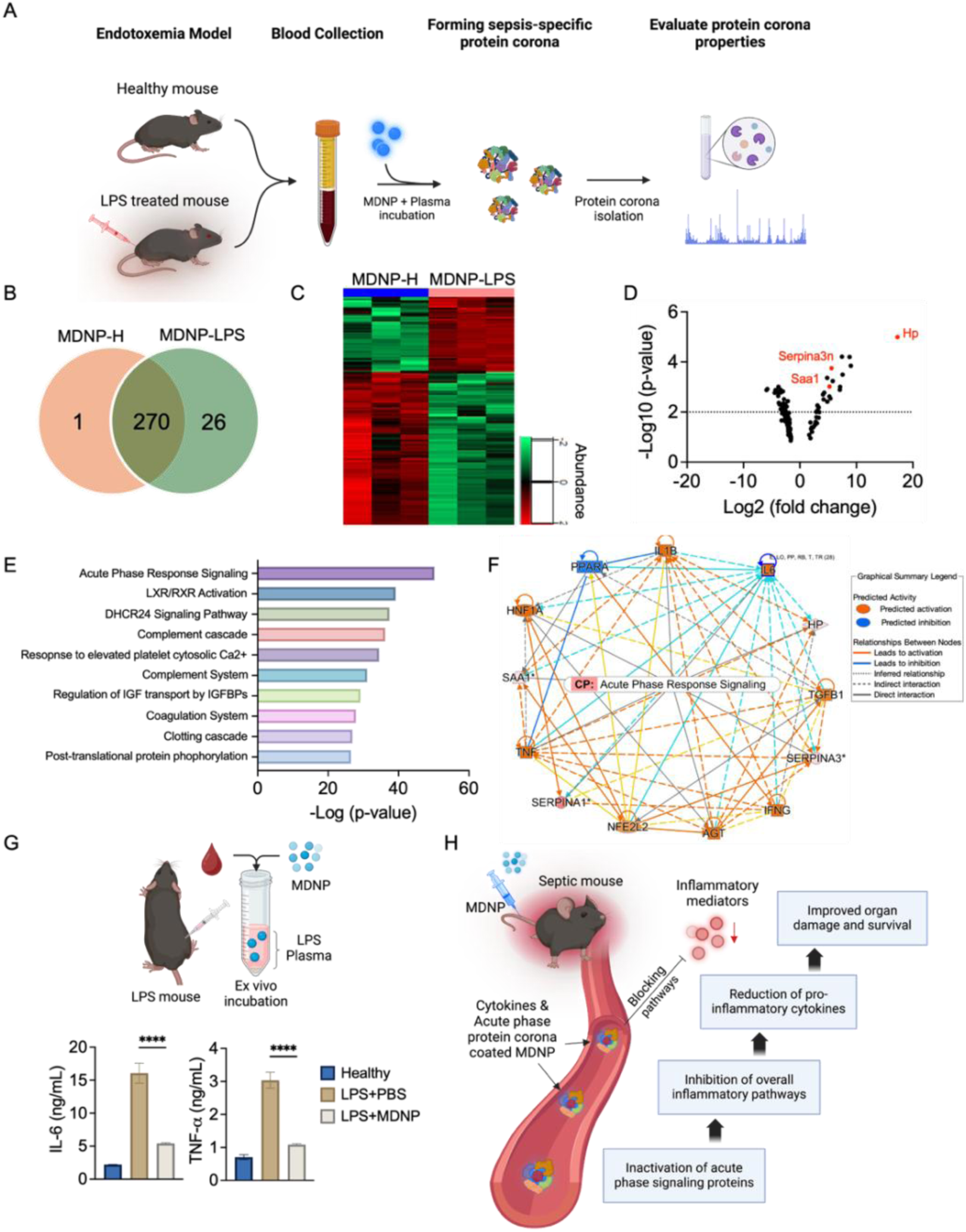
Mapping and identification of protein corona adsorbed onto the surface of MDNP. (A) A schematic showing the process of evaluating the MDNP protein corona. (B) Venn diagram illustrating a total of 296 proteins detected including unique (1 in healthy and 26 in LPS) and 270 shared proteins in the coronas. (C) Heatmap shows abundance of 49 upregulated and 100 downregulated proteins in MDNP-LPS group compared to MDNP-H healthy group. (D) Volcano plot of significantly upregulated and downregulated corona proteins. Hp, Serpina3n and Saa1 (red dots) are categorized as pro-inflammatory and acute phase response signaling proteins. (E) Top 10 canonical pathways associated with the significantly upregulated protein corona were identified, with acute phase response signaling being the most prominent pathway. (F) A network representation of acute phase response signaling proteins and major upstream regulators. (G) *Ex vivo* sequestration of pro-inflammatory cytokines by incubating LPS plasma with MDNP. Both IL-6 and TNF-α was significantly reduced by 8 hr treatment. (H) A graphical summary illustrating how the cytokine and acute phase protein corona-coated MDNP sequesters and neutralizes inflammatory proteins, thereby reducing overall inflammation.

**Table 1.**
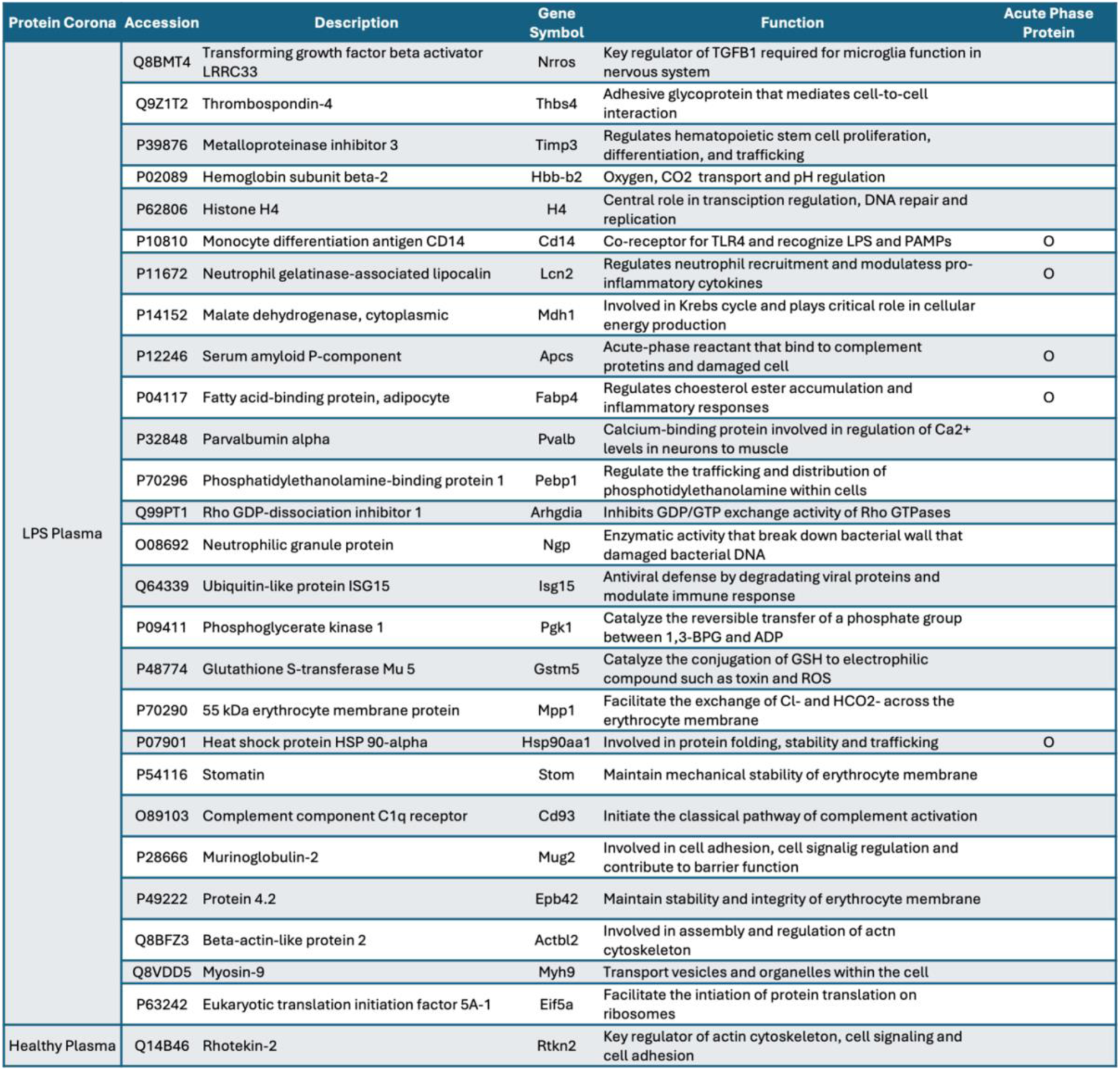
A list of unique proteins that are part of the MDNP protein corona formed from LPS and healthy plasma. A total 26 proteins are derived from the LPS plasma protein corona, while only one unique protein is found in healthy plasma corona. Acute phase proteins are highlighted with circles to underscore the potential presence of pro-inflammatory proteins associated with MDNP.

### MDNP treatment improved survival in polymicrobial sepsis model

The survival benefit of MDNP was also tested in a clinically relevant severe polymicrobial cecal ligation and puncture (CLP) sepsis model (**Figure 7A**). Mice underwent the CLP procedure and were treated with three doses of 2 mg MDNP, followed by monitoring for 5 days. To confirm MDNP could sequester pro-inflammatory cytokines in CLP plasma, we first incubated MDNP with plasma isolated at 3 hr post-CLP and observed an almost complete reduction in IL-6 and TNF-α (**Figure 7B**). In the survival study, MDNP treated mice showed significant improvement in survival (40%) compared to control group (0%) (**Figure 7C**). Taken together, the ability of MDNP to increase the survival of septic mice in the absence of antibiotics, highlights their translational potential for treating inflammatory diseases, including sepsis.

**Figure 7.**
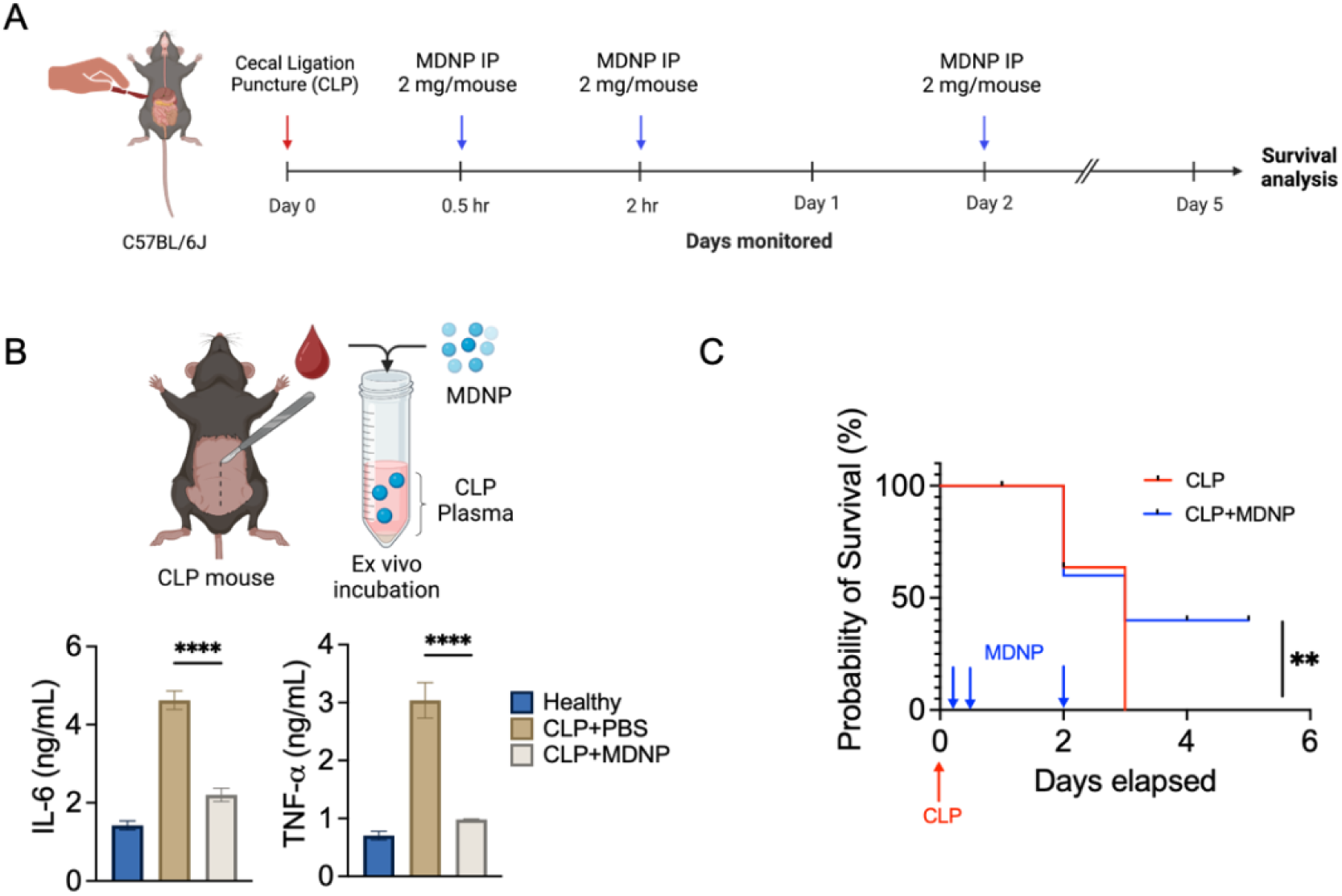
Survival effects of MDNP treatment on severe CLP model. (A) Schematic timeline of survival study of cecal ligation and puncture (CLP) C57BL/6J murine model with MDNP treatment (n=5/group) based on the complement cascade pathway from proteomics. Mice underwent CLP surgery to induce severe polymicrobial inflammation and three separate doses of MDNP were administered intraperitoneally (0.5, 2, 24 hr timepoint, 2 mg/injection) to analyze survival. (B) *Ex vivo* sequestration test of CLP plasma and MDNP showed that both IL-6 and TNF-α was significantly reduced. Kaplan-Meier curves and a log-rank test was performed to compare the survival rates. (C) The survival test displayed a significant improvement (40%) by the treatment of MDNP **p< 0.005. Data are expressed as mean ± SD (n=5). ****p<0.0005 versus PBS control group.

## Discussion

A tremendous amount of research has been implemented to enhance the clinical management of inflammatory diseases caused by excessive stress (Miller and Raison, 2016), infections(Ikoba et al., 2015), autoimmune disorders (Miller et al., 2007), or secondary diseases (Dufour et al., 2022). Despite many clinical trials aimed at blocking specific inflammatory mediators, most have been unsuccessful due to the complexity and multifactorial nature of inflammatory diseases. This highlights the need for multimodal approaches that target multiple inflammatory mediators simultaneously to improve treatment outcomes. The core pathophysiology of sepsis is known to involve the disruption of the finely tuned balance between inflammatory and anti-inflammatory immune responses (Jarczak et al., 2021). Recognition of pathogen- or damage-associated molecular patterns (PAMPs/DAMPs) by innate immune cells functions as the trigger to initiate inflammatory cell signaling and subsequent cytokine release, leading to multiorgan damage, and death. Notably, studies have shown that increased plasma IL-6 and TNF-α levels in sepsis patients are correlated with mortality (Mera et al., 2011; Unsinger et al., 2010; Wu et al., 2009). Our MDNP efficiently reduced systemic pro-inflammatory cytokines including IL-6, TNF-α, among others (**Figure 4E, 6G, 7B**), while simultaneously neutralizing additional pro-inflammatory mediators (APP) (**Figure 6D**).

Strategies for reducing pro-inflammatory mediators include conventional adsorption resins like Cytosorb®, or more sophisticated membrane-coated nanoparticles (Thamphiwatana et al., 2017), telodendrimer nanotraps (Shi et al., 2020), or abiotic histone-capturing hydrogel nanoparticles (Koide et al., 2021b). Although these approaches have shown success in certain cases, the complex structural composition and manufacturing processes may hinder rapid development. In contrast, we developed a MDNP, isolated from maca root, as a novel approach to mitigate broad pro-inflammatory responses from abundant and readily available plant material, facilitating convenient access, and long-term storage potential (**Figure 1**). We further demonstrated their ability to reduce organ damage and improve survival as reflected in two representative and lethal sepsis models (**Figure 5A, 5H, 7C**) through the formation of a multimodal protein corona (**Figure 6**).

Sepsis endotypes include hyperinflammatory (early) and immunosuppressive (late) states (Hotchkiss et al., 2013). A clinical study indicated that early-identified sepsis is associated clinically with higher mortality than late-identified sepsis (Jee et al., 2020). Thus, achieving control over the cytokine storm holds great potential to improve patient survival. Modes of cytokine sequestration have generally relied on charge-based interactions, where most pro-inflammatory cytokines are negatively charged, and anti-inflammatory cytokines are positively charged at neutral pH (Messina et al., 2024).

Interestingly, the zeta potential of MDNP was −9.6 mV (**Figure 1E**), yet cytokines like IL-6 (pI 6.96) and TNF-α (pI 5.01) were strongly sequestered but IL-1β (pI 4.96) and IFN-γ (pI 8.25) were not (**Figure 2K, L, S6**). These differences suggest the possibility of a combination of forces driving the binding of cytokines to the surface of MDNP. Our lipidomics analysis of MDNP (**Figure 1H, I**) showed the presence of multiple charged or ionizable lipid species like Cer (Phyto), HexCer (Phyto), FFA, LPC, and PC (**Supplementary Figure S2**). Therefore, it is possible that a combination of local charge and hydrophobic interactions are contributing to the apparently selective sequestering of specific pro-inflammatory cytokines. Although we have not fully elaborated how MDNP facilitate a selective binding interaction for certain cytokines, which is a limitation of our study, our results provide the basis for future studies to investigate the role of specific lipid components and their ability to alter binding interactions.

Upon exposure to biological fluids, nanoparticles dynamically adsorb biomolecules onto their surface, forming a protein corona that dictates how nanoparticles interact with host cells. These interactions can alter both drug delivery efficiency (Corbo et al., 2016) and biological function (Cedervall et al., 2007) of the particles. The analysis of MDNP protein coronas revealed significant differences between MDNP-H and MDNP-LPS groups, highlighting that unique protein fingerprints are associated with MDNP under inflammatory conditions. This is consistent with the concept of the ‘personalized protein corona’, which demonstrates that unique protein fingerprints are formed on the surface of nanoparticles as a function of disease state (Ju et al., 2020; Shaw and Pearson, 2022). The ability of MDNP to bind cytokines in LPS- or CLP-treated mouse plasma confirmed its ability to directly modulate inflammatory responses in a disease setting (**Figure 6G, 7B**). An unexpected finding from our proteomics analysis was the identification of multiple pro-inflammatory APP being upregulated in the coronas of MDNP-LPS like Hp, SerpinA3N, and Saa1 (**Figure 6D**). Hp activates immune cells, promotes secretion of pro-inflammatory cytokines, and interacts with complement system, potentially intensifying inflammatory responses (Torres et al., 2022). SerpinA3N is involved in regulating inflammatory responses by controlling protease activity to prevent excessive tissue damage, but it’s upregulated under inflammatory conditions (Liu et al., 2023). Saa1 plays direct role in recruiting immune cells and modulating pro-inflammatory cytokines (Ye and Sun, 2015). Supporting that MDNP neutralized the pro-inflammatory functions of corona constituents, MDNP-LPS was cultured with BMDM and no significant increase in IL-6 or TNF-α was detected (**Supplementary Figure S9**). These results highlighted that MDNP formed multifaceted protein corona under inflammatory conditions that could mitigate aberrant immune activation caused by individual components, representing a novel approach to the treatment of inflammatory diseases.

Here, we demonstrated that MDNP sequestered and neutralized multiple pro-inflammatory cytokines and APP in its protein corona to reveal a highly effective anti-inflammatory plant-derived therapeutic strategy for managing severe inflammatory responses. Our data indicates that MDNP exhibit negligible toxicity, are efficiently internalized by macrophages, display immunomodulatory activities, and demonstrate significant therapeutic efficacy. Administration of MDNP *via* IP or IV routes effectively reduced pro-inflammatory cytokines and promoted improved survival in two representative sepsis mouse models, LPS-induced endotoxemia and CLP polymicrobial sepsis. This research underscores the broad therapeutic potential of the MDNP, which leverages a unique and never previously examined mechanism of action through the formation of a multimodal protein corona that effectively binds and neutralizes pro-inflammatory cytokines and APP from propagating systemic inflammation.

## Materials and methods

### Isolation, purification, and characterization of maca-derived lipid nanoparticles (MDNP)

A fine ground organic powder of 100% *Lepidium meyenii* Walp (maca) (Family: Brassicaceae, Order: Brassicas) was purchased from an organic producer, Happy Andes (https://www.happyandes-usa.com/). Maca powder (30 g) was mixed in 400 mL of autoclaved deionized water and stirred for 12 hrs at room temperature to make maca juice. The obtained maca juice was transferred to 50 mL tubes and centrifuged at 3,000 x g for 20 min at 20°C then the supernatant was centrifuged again in new 50 mL tubes at 10,000 x g for 2 hrs at 20°C to remove small and large fibers. Next, the clear light brown colored supernatant was collected in 70 mL polycarbonate ultracentrifuge bottles (Beckman Coulter, Brea, CA) and was ultracentrifuged with using a 45 Ti rotor (Beckman Coulter, Brea, CA) at 150,000 x g for 2 hrs at 4°C. Then, the collected pellets were suspended in 2 mL of phosphate-buffered saline (PBS) through dispersion with an ultrasonic processor. PBS-suspended maca pellets were transferred to a sucrose gradient (8%, 20%, 35% [g/v]) and ultracentrifuged at 150,000 x g for 2 hrs at 4°C, then the cloudy band between 8% and 20% sucrose was collected as previously described (Sung et al., 2019). The concentration of MDNP was measured using a Bio-Rad quantification assay. The quantified MDNP were stored at 4°C. The size and zeta potential of MDNP was measured by dynamic light scattering (DLS) using a Zetasizer Nano ZSP (Malvern, UK). MDNP were also tested under different storage temperature conditions (room temperature (RT), 4°C, −20°C, and −80°C) to determine their stability. Transmission electron microscopy (TEM) images of MDNP were acquired using a formvar-coated copper grid and staining using 1% uranyl acetate. Differential Scanning Calorimeter (DSC) 2500 (TA Instruments, New Castle, DE) was used to analyze the temperature and heat flow associated with thermal transition in the MDNP. 5 µL of sample in liquid form was added into a hermetic aluminum pan and sealed with metal lid (DSC Consumables, Austin, MN) with Tzero sample press kit (TA Instruments, New Castle, DE). The following thermal procedure was used to scan: ramp 2°C/min from 25°C to 80°C. The heat flow transition was analyzed on a plot by TA analysis software Trios (Version 5.4.0.300). For lyophilization, frozen MDNP in 1.5 mL tubes with holes on top of screw caps were placed in a benchtop freeze-dryer (Labconco, Kansas City, MO) overnight. Completely dried samples were collected and reconstituted in PBS for stability testing.

### Lipid extraction and composition analysis

A total lipid extract was prepared using a modified methyl-tert-butyl ether (MTBE) lipid extraction protocol (Matyash et al., 2008). Briefly, 400 µL of cold methanol and 10 µL of internal standard mixture (EquiSPLASH lipidomix) were added to each sample followed by incubation at 4°C, 650 RPM shaking for 15 min. Next, 500 µL of cold MTBE was added followed by incubation at 4°C for 1 hr with 650 RPM shaking. 500 µL of cold water was added slowly and resulting extract was maintained 4°C, 650 rpm shaking for 15 min. Phase separation was completed by centrifugation at 8,000 RPM for 8 min at 4°C. The upper, organic phase was removed and set aside on ice. The bottom, aqueous phase was re-extracted with 200 µL of MTBE followed by 15 min of incubation at 4°C with 650 RPM shaking. Phase separation was completed by centrifugation at 8,000 RPM for 8 min at 4°C. The organic extract was dried under a steady stream of nitrogen at 30°C. The recovered lipids were reconstituted in 200 µL of chloroform:methanol (1:1, v/v) containing 200 µM of butylated hydroxytoluene. Prior to analysis, samples were further diluted with acetonitrile:isopropanol:water (1:2:1, v/v/v). The total lipid extract was analyzed by liquid chromatography coupled to high resolution tandem mass spectrometry (LC-MS/MS). The LC-MS/MS analyses were performed on an Agilent 1290 Infinity LC coupled to an Agilent 6560 Quadrupole Time-of-Flight (Q-TOF) mass spectrometer. The separation was achieved using a C18 CSH (1.7 µm; 2.1 x 100 mm) column (Waters, Milford, MA). Mobile phase A was 10 mM ammonium formate with 0.1% formic acid in water/acetonitrile (40:60, v/v) and mobile phase B was 10 mM ammonium formate with 0.1% formic acid in acetonitrile/isopropanol (10:90, v/v). The gradient was ramped from 40% to 43% B in 1 min, ramped to 50% in 0.1 min, ramped to 54% B in 4.9 min, ramped to 70% in 0.1 min, and ramped to 99% B in 2.9 min. The gradient was returned to initial conditions in 0.5 min and held for 1.6 min for column equilibration. The flow rate was 0.4 mL/min. The column was maintained at 55°C and the auto-sampler was kept at 5°C. A 2 µL injection was used for all samples. LC-MS data from the iterative MS/MS workflow was analyzed for lipid identification via Agilent’s Lipid Annotator (version 1.0). Positive and negative ion mode adducts included [M+H]^+^, [M+Na]^+^, [M+NH_4_]^+^, [M-H]^-^, and [M+CH_3_CO_2_]^-^, respectively. The LC-MS data from the MS^1^ workflow were processed using Agilent’s MassHunter Profinder (version 10.0).

### Characterization of MDNP protein composition and MDNP protein corona fingerprints in healthy and LPS plasma

To form the MDNP protein corona, purified MDNP were incubated with plasma isolated from healthy and LPS-treated C57BL/6J mice for 4 hrs at 37°C. MDNP bound to proteins were washed and centrifuged 3 times at 13,000 xg, 4°C with 1x cold PBS. Then it was evaluated using the below protocol. Prepared samples were lysed in a lysis buffer containing 5% sodium dodecyl sulfate (Sigma, St. Louis, MO), 50 mM triethylammonium bicarbonate (1M, pH 8.0) (Sigma, 7408). Proteins were extracted and digested using S-trap micro columns (ProtiFi, Farmingdale, NY). The eluted peptides from the S-trap column were dried, and peptide concentration was determined using a BCA assay kit (Thermo Fisher Scientific, 23275), after reconstitution in 0.1% formic acid. All tryptic peptides were separated on a nanoACQUITY Ultra-Performance Liquid Chromatography analytical column (BEH130 C18, 1.7 µm, 75 µm x 200 mm; Waters Corporation, Milford, MA, USA) over a 185-min linear acetonitrile gradient with 0.1% formic acid on a nanoACQUITY Ultra-Performance Liquid Chromatography system (Waters Corp, Milford, MA) and analyzed on a coupled Orbitrap Fusion Lumos Tribrid mass spectrometer (Thermo Scientific, San Jose, CA). Full scans were acquired at 240,000 resolutions, and precursors were fragmented by high-energy collisional dissociation at 35% for up to 3 sec. MS/MS raw files were processed using Thermo Proteome Discoverer (PD, version 3.0.0.757) with the Sequest HT search engine against the plant metagenome database and UniProt mouse reference proteome. Trypsin was used with a maximum of two missed cleavages, and peptide lengths were restricted to 6-144 amino acids. Label-free quantification was performed using the Minora feature detector. Protein identification was filtered at a 1% false discovery rate across PSM, peptide, and protein levels using the Percolator algorithm in PD. Exported protein abundances were analyzed with Perseus software (version 1.6.14.0), with further filtering to include only proteins identified without missing values across all samples. The quantitative protein data were log_2_ transformed and further normalized using median centering. Two-tailed student’s t-test (adjusted p-value < 0.05) was applied to determine the differentially expressed proteins (DEPs). Plant data was analyzed with library from Uniprot and the Plant Proteome Database (Lepidium meyenii: taxonomy ID 153348, entry: 92 sequences; Plant metagenome: taxonomy ID 1297885, entry: 27312 sequences; Zingiberales: taxonomy ID 4618, entry: 357900) Ingenuity Pathway Analysis (Qiagen) was utilized to identify canonical pathways, biological function, upstream regulators and disease association.

### Cell culture

The generation of bone marrow-derived macrophages (BMDM) followed a previously published method(Truong et al., 2024). As established, the tibia and femur from a C57BL/6J mouse was isolated by removing bulk muscles and connective tissues. RPMI media (supplemented with L-glutamine, penicillin (100 units/mL), streptomycin (100 µg/mL), 10% heat-inactivated FBS, and 20% L929 cell-conditioned media) drawn needles were inserted into bones to flush the marrow into a 10 cm petri dish. The collected bone marrow was filtered to grow in uncoated 10 cm cell culture plates. BMDM were cultured in RPMI media conditioned with L929 at 37°C, 5% CO_2._ The media was replaced every 3 days. On Day 8-10, BMDM were lifted using Versene (Gibco, Grand Island, NY), to be used for subsequent experiments. Trypan blue solution was used to determine the cell number and viability with an EVE^TM^ Automated Cell Counter (NanoEntek, Waltham, MA). RAW-blue cells were also cultured to confluency in 75 cm^2^ flasks in identical incubation condition with Dulbecco’s Eagle Medium (DMEM) supplemented with penicillin (100 U/mL), streptomycin (100 U/mL), and heat inactivated fetal bovine serum (10%).

### Mice

Male C57BL/6 (6 to 8-week-old) purchased from the Jackson Laboratory (Bar Harbor, ME) were maintained in cages at ambient temperature, 55% relative humidity, and under a 12 hr dark/light cycle. LPS-induced endotoxemia model: Mice were challenged with 5 mg/kg LPS intraperitoneally (IP) prior to treatment with two doses of 2 mg MDNP at 30 min and 2 hr *via* both IP and intravenous (IV) injection. Cecal ligation and punction (CLP) model: Mice were lightly anesthetized with a mixture of ketamine (75 mg/kg) and xylazine (15 mg/kg) at 1:1 ratio. After abdominal fur was removed, a small incision was made to expose the cecum. The cecum was then ligated with a silk suture and perforated with 19-gauge needle. A small amount of feces was extruded by gently squeezing the cecum, the cecum was replaced, and the abdomen was sutured after the bowel was repositioned. Experimental groups were IP injected with three doses of 2 mg MDNP at 30 min, 2 hr, and 24 hr timepoint. All experiments were performed in compliance with the protocol by the University of Maryland, Baltimore Institutional Animal Care and Use Committee (IACUC) as well as the ARRIVE guidelines.

### Cytotoxicity assay

To measure the cytotoxicity of MDNP, a PI/Annexin V-FITC apoptosis assay was conducted using BMDM. The assay utilizes propidium iodide (PI) staining to distinguish between early and late-stage apoptotic cells. FITC labeling allow for the visualization of Annexin V-bound cells using flow cytometry. Cultured BMDM at a density of 1.0 x 10^5^ per well were treated with MDNP at 1 mg/mL for 8 hrs, then the cells were harvested with Versene after washing with PBS. Collected cells were centrifuged at 500 x g for 5 min and cell pellets were resuspended in 1x binding buffer. 100 μL of cell suspension in flow cytometry tubes were added with Annexin V-FITC and PI to be analyzed by flow cytometry. The data was analyzed by De Novo software FCS Express 7 (Dotmatics, Boston, MA).

### *In vitro* cellular internalization

BMDM were seeded overnight in Falcon Culture slides at a density of 0.5 x 10^5^ per well. Cy5.5 was incubated with MDNP under shaking for 30 min in the dark to label the MDNP and excessive Cy5.5 dye was removed by 20 min centrifugation at 13,000 x g prior to use. Then, 50 μL of Cy5.5-labled MDNP were added to 0.5 mL of culture medium and incubated with cells for 0, 4, 8 hrs. Negative control, untreated cells were used to establish the 0 hr timepoint. At the determined timepoints, cells were washed with PBS three times, fixed with 4% paraformaldehyde (PFA) for 10 min, and dehydrated with acetone for 5 min at −20°C. After blocking the culture with 1% BSA in 1x PBS 30 min, the cells were washed again with 1x PBS and treated with 100 µL of fluorescein isothiocyanate (FITC)-labeled phalloidin (1:50 dilution in PBS) for 30 min to stain F-actin. The cells were then washed two times with 1x PBS and dried in the dark condition and glass cover slips were mounted with mounting medium containing 4’,6-diamidino-2-phenylindole (DAPI) after carefully removing the plastic chambers. The final fluorescence images were captured using an Olympus fluorescence microscope (Tokyo, Japan) equipped with Hamamatsu Digital Camera ORCA-03G and Nikon software was used to analyze the image data.

### Flow cytometric phenotyping of MDNP-treated macrophages

BMDM were seeded at a density of 1.0 x 10^5^ per well were in a sterile 24 well plate. Cells were treated with 100 ng/mL of LPS in complete media for 24 hrs to induce inflammation. Then, 100 μg/mL and 200 μg/mL MDNP was treated for 8 hrs. Cells were resuspended in MACS buffer (PBS pH 7.2 supplemented with 1% FBS and 0.4% 0.5 M EDTA, Quality Biological, Gaithersburg, MD) and transferred to the flow cytometry tubes. FcR blocking (CD16/32, Biolegend, San Diego, CA) was performed and cells were stained with following surface marker antibodies: Live/Dead fixable green, F4/80, CD11b, MHCII, CD206, CD80, and CD86 (Biolegend, San Diego, CA). Samples were analyzed using Cytek Aurora 3 (Fremont, CA). FCS Expression 7 Flow Cytometry De Novo Software was used for flow data processing. Mean fluorescence intensity (MFI) was generated by GraphPad Prism Software 10.1.1.

### *In vitro* pro-inflammatory cytokine sequestration and effects on NF-κB activity

Anti-inflammatory properties of MDNP were investigated in two ways; prophylactic and therapeutic. For prophylactic treatment, BMDM at a density of 1.0 x 10^5^ per well were treated with 100 μg/mL MDNP for 8 hrs and 100 ng/mL of LPS in complete media in 24 well plates (Corning, Corning, NY) were incubated for 4 hrs. For therapeutic study, BMDM in identical culture condition as above were treated with 100 ng/mL of LPS for 4 hrs and 100 µg/mL of MDNP was subsequently added and incubated for 8 hrs. Supernatants were then collected to measure the reduction of pro-inflammatory cytokines including IL-6, TNF-α, IL-1β, and IFN-γ by enzyme-linked immunosorbent assay (ELISA) (BioLegend, San Diego, CA). We also assessed the ability of MDNP to modulate cytokine levels in the absence of cells. The supernatants from another set of LPS-treated BMDM were collected and incubated with 100 µg/mL of MDNP in a time dependent manner (0, 0.5, 1, 4, 8, 12, and 24 hrs). Lastly, we confirmed the direct ability of MDNP to sequester IL-6 by incubating 5 ng/mL of IL-6 recombinant protein in DMEM with 100 µg/mL of MDNP for 8 hrs. Remaining IL-6 levels were measured using ELISA.

An NF-*κ*B reporter cell line, RAW-Blue, was used to determine the ability of MDNP to alter NF-*κ*B activity due to LPS stimulation. The cells were plated at a density of 1.0 x 10^5^ per well and were stimulated with LPS for 4 hrs and treated with 100 µg/mL MDNP for 8 hrs. Then, QUANTI-Blue was added to the collected supernatant in flat bottom 96 well plate. It was incubated at 37°C for 1 hr, then secreted embryonic alkaline phosphatase (SEAP) reporter was detected using a spectrophotometer at 620-655 nm. Poly(lactic-co-glycolic acid) (PLGA) particles of similar size as MDNP were used as a control and fabricated using a microfluidics device as we previously described(Truong et al., 2021). The size and zeta potential of the PLGA was measured 163.8 nm and −39.3 mV, respectively.

### *Ex vivo* cytokine sequestration assay

To evaluate the cytokine sequestration ability of MDNP in mouse plasma *ex vivo*, 100 µg/mL of MDNP in PBS was prepared and added to 100 µL of diluted LPS-treated or CLP mouse plasma to ultracentrifuge tubes. The mixture tubes were incubated at 37°C for 8 hrs. After incubation, samples were centrifuged at 2,000 x g for 10 min to pellet the cytokine bound MDNP. Then the supernatant was collected to measure IL-6 and TNF-α levels using ELISA (Biolegend, San Diego, CA).

### *In vivo* biodistribution and anti-inflammatory effect following LPS challenge

The *in vivo* biodistribution study was performed by intraperitoneal (IP) or intravenous (IV) injection of MDNP that were labeled with the near-infrared fluorescent Cy5.5 dye. Mice were administered 2 mg of Cy5.5-MDNP twice (0.5 and 2 hr time point)-post LPS challenge (5 mg/kg) and euthanized after 4 hrs to collect blood by cardiac puncture and organs including liver, lung, spleen, heart, and kidneys. The fluorescence of the organs was imaged using an *in vivo* imaging system (IVIS) with emission (720 nm) upon laser excitation (675 nm). Blood samples were centrifuged at 1,000 x g for 20 min at 4°C in microtainer capillary blood collection plasma tubes. The plasma samples were used to measure the alteration of systematic cytokines levels using a Luminex MAGPIX System (Luminex, Austin, TX). The 7-plex panel included murine IL-6, TNF-α, MCP-1, IL-10, IL-1β, IFN-β, and IFN-γ. Blood biochemistry was also measured for alanine aminotransferase (ATL/SGPT), aspartate aminotransferase (AST/SGOT), creatine (CREA), blood urea nitrogen (BUN), total protein (TP), and globulin, and albumin (VRL Animal Health Diagnostics, Gaithersburg, MD). The collected organs were fixed in 4% formalin immediately after euthanasia and paraffin-embedded for sectioning. Hematoxylin & eosin (H&E) staining was performed using standard procedures by the Pathology Biorepository Shared Services Core at the University of Maryland, Baltimore. The level of injury scores was calculated based on the scoring criteria: score 0, no damage; score 1, >10% slight inflammation; score 2, 10-25% mild; score 3, 26-50% moderate; score 4, 51-75% severe; score 5, >75% necrosis.

### *In vivo* sepsis survival study

The *in vivo* survival study was carried out using two representative sepsis models: a lethal LPS-induced endotoxemia and CLP polymicrobial mouse model. For endotoxemia, C57BL/6J mice were challenged with 20 mg/kg LPS and two doses MDNP (2 mg) administered *via* IP injection. For CLP model, the cecum of anesthetized C57BL/6J mice was perforated to release fecal materials into peritoneal cavity to generate an exacerbated polymicrobial infection. Three doses of MDNP (2 mg/injection) were administered via IP route. Survival for each model was evaluated over the course of 5 days. Body temperature was also recorded every day for CLP mice.

### Statistical analysis

Data evaluation was performed using GraphPad Prism Software 10.1.1 (San Diego, CA). One-way ANOVA and t-test for unpaired data were employed to evaluate statistical significance along with Tukey’s multiple comparison test. Kaplan-Meier survival curve and statistical significance of mouse survival were determined with a long-rank (Mantel-Cox) X^2^ test. Significant differences were indicated as p<0.05, p<0.005, p<0.0005 related to control unless otherwise stated.

## Abbreviations

AP-1: activator protein-1
APCS: Serum Amyloid P component
APP: acute phase proteins
APR: acute phase response
AST/SGOT: aspartate aminotransferase
ATL/SGPT: alanine aminotransferase
BMDM: bone marrow derived macrophages
BUN: blood urea nitrogen
Cer(Phyto): phytoceramide
CLP: cecal ligation and puncture
CREA: creatine
DAMP: damage-associated molecular pattern
DAPI: 4’,6-diamidino-2-phenylindole
DGDG: digalactosyldiacylglycerol
DLS: dynamic light scattering
DSC: differential scanning calorimetry
FABP4: Fatty Acid Binding Protein 4
FFA: free fatty acids
FITC: fluorescein isothiocyanate
H&E: hematoxylin & eosin
HexCer(phyto): phytohexosylceramide
Hp: haptoglobin
HSP90AA: Heat Shock Protein 90 alpha
IFN-γ: interferon-gamma
IL-1β: interleukin-1beta
IL-6: interleukin-6
IP: intraperitoneal
IV: intravenous
LCN2: lipocalin-2
LPC: lysophosphatidylcholine
LPS: lipopolysaccharide
MDNP: maca-derived lipid nanoparticles
NF-*κ*B: nuclear factor kappa-light-chain-enhancer of activated B cells
NGP4: neutrophilic granule protein
PAMP: pathogen-associated molecular pattern
PC: phosphatidylcholine
PDNP: plant-derived lipid nanoparticle
PEG: poly(ethylene glycol)
PI: phosphatidylinositol
PLGA: poly(lactic-co-glycolic acid)
Saa1: serum amyloid A1
SEAP: secreted alkaline phosphatase
SerpinA3N: serine protease inhibitor A3N
TG: triglyceride
TLR-4: toll-like receptor-4
TNF-α: tumor necrosis factor-alpha
TP: total protein

## Data availability statement

All lipidomics and proteomics raw data are included in the supporting information. Additional datasets supporting the findings are available upon reasonable request from the corresponding author.

## Acknowledgements

This work was supported by Startup funds from the University of Maryland School of Pharmacy, the National Institute of General Medical Sciences of the National Institute of Health under Award number R35GM142752 awarded to R.M.P. A.L.C. was supported by a pharmaceutical sciences (PSC) departmental fellowship. This publication was supported by funds through the Maryland Department of Health’s Cigarette Restitution Fund Program and the National Cancer Institute – Cancer Center Support Grant (CCSG) – P30CA134274. University of Maryland School of Medicine’s & Greenebaum Comprehensive Cancer Center’s Flow Cytometry Core–Baltimore, Maryland. University of Maryland School of Medicine’s Center for Translational Research in Imaging – Baltimore, Maryland. University of Maryland School of Pharmacy Mass Spectrometry Center (SOP1841-IQB2014). BioRender was used for preparing graphics. The content is solely the responsibility of the authors and does not necessarily represent the official view of the National Institutes of Health.

## Author contributions

**Junsik J. Sung:** Conceptualization, Formal analysis, Investigation, Methodology, Visualization, Writing – original draft, Writing – review & editing. **Jacob R. Shaw:** Methodology, Writing – review & editing. **Josie D. Rezende:** Investigation, Methodology. **Shruti Dharmaraj:** Investigation, Methodology. **Andrea L. Cottingham:** Investigation, Methodology. **Mehari M. Weldemariam:** Investigation, Methodology. **Jace W. Jones:** Investigation, Methodology, Resources, Supervision. **Maureen A. Kane:** Investigation, Methodology, Resources, Supervision. **Ryan M. Pearson:** Conceptualization, Funding acquisition, Methodology, Resources, Supervision, Writing – original draft, Writing – review & editing.

