## Supplemental data for "Lipid Nanoparticles from *L. meyenii* Walp Mitigate Sepsis through Multimodal Protein Corona Formation"

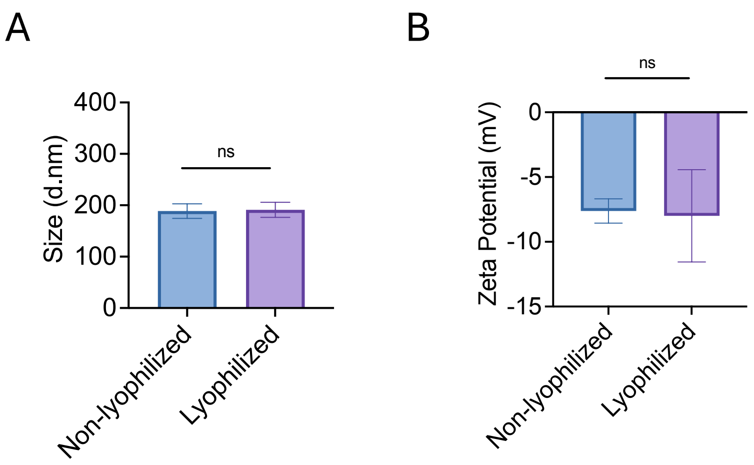


**Supplementary Figure S1**. Stability test for lyophilized MDNP (A) Size was stable after reconstituting lyophilized MDNP (B) Zeta potential of lyophilized MDNP also displayed stable measurement.

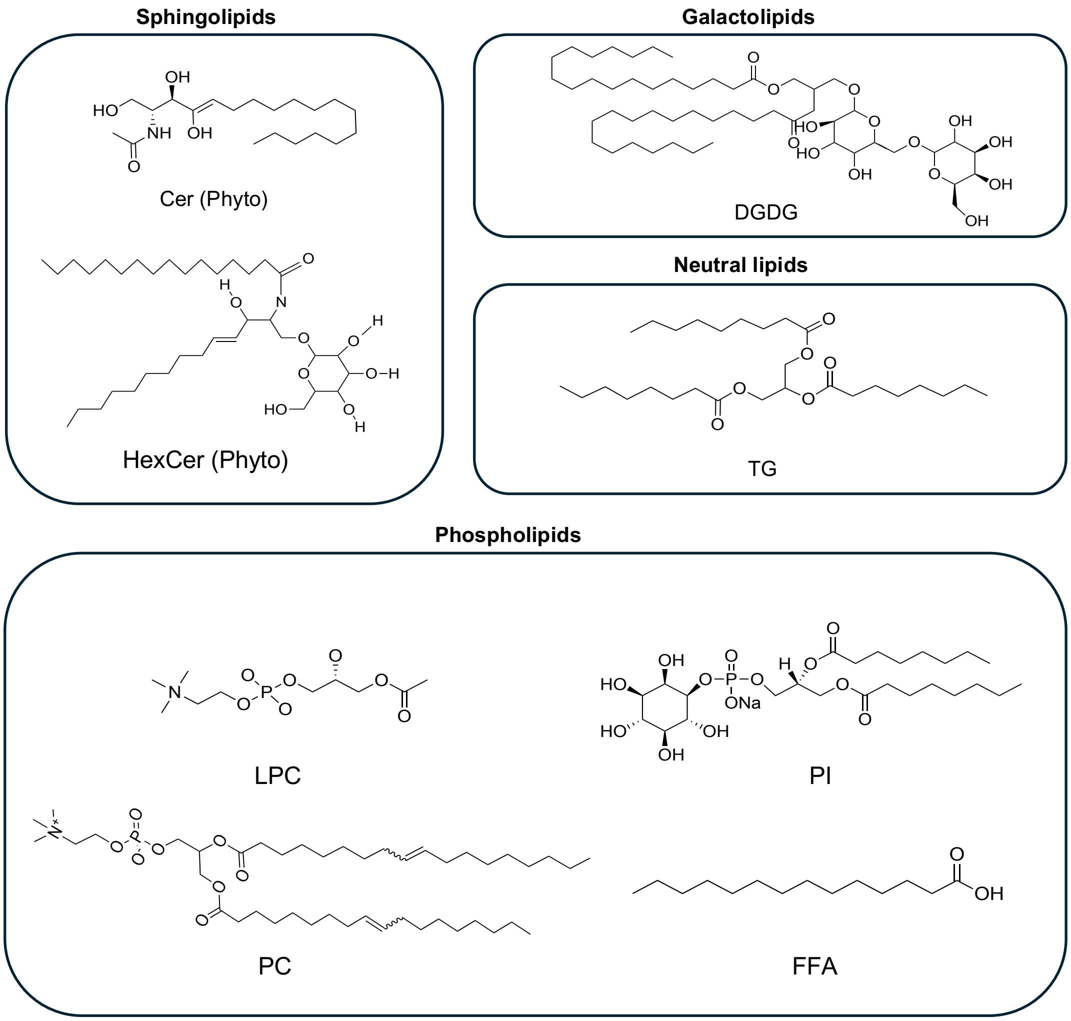


**Supplementary Figure S2.** Chemical structure of lipid components in four distinctive lipid groups (sphingolipids, galactolipids, neutral, and phospholipids).

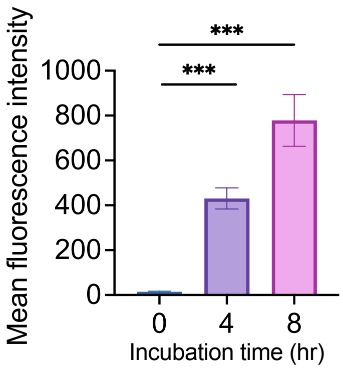


**Supplementary Figure S3**. Quantitative measure of mean fluorescence intensity (MFI) from the uptake of Cy5.5-labeled MDNP at 0, 4, 8 hr time point. A significant uptake was detected by confocal microscopy over time.

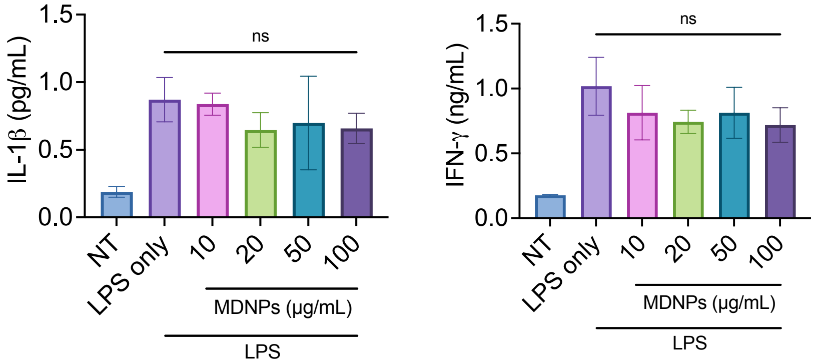


**Supplementary Figure S4.** Therapeutic effect of MDNP on pro-inflammatory cytokines *in vitro*. Unlike IL-6 and TNF-α, IL1β and IFN-γ was not affected by the treatment.

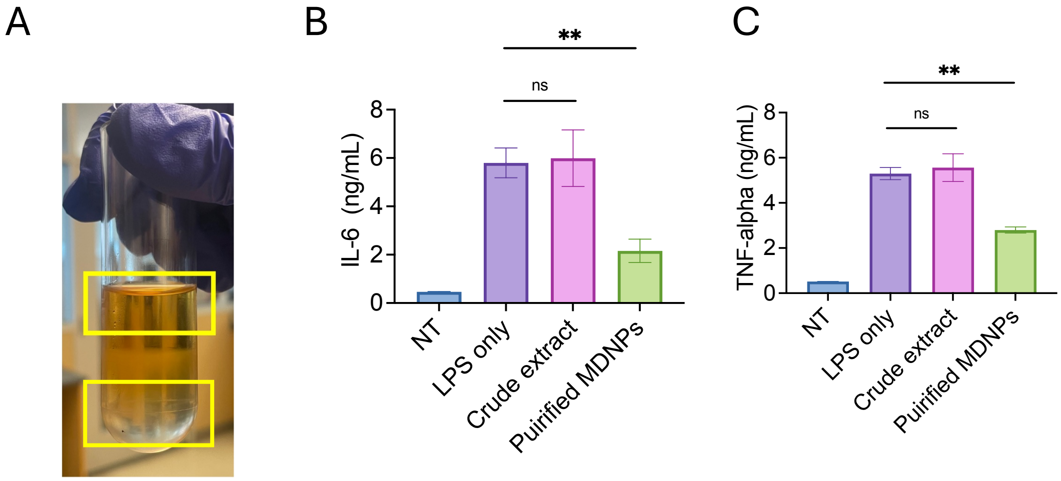


**Supplementary Figure S5.** Incapability of anti-inflammatory effect in crude extract after MDNP isolation (A) Crude extract in yellow rectangles is the remaining solution after MDNP isolation through sucrose gradient separation. Result does not display significant removal effect on pro-inflammatory cytokines (B) IL-6 and (C) TNF-α by treatment of crude extract from maca juice.

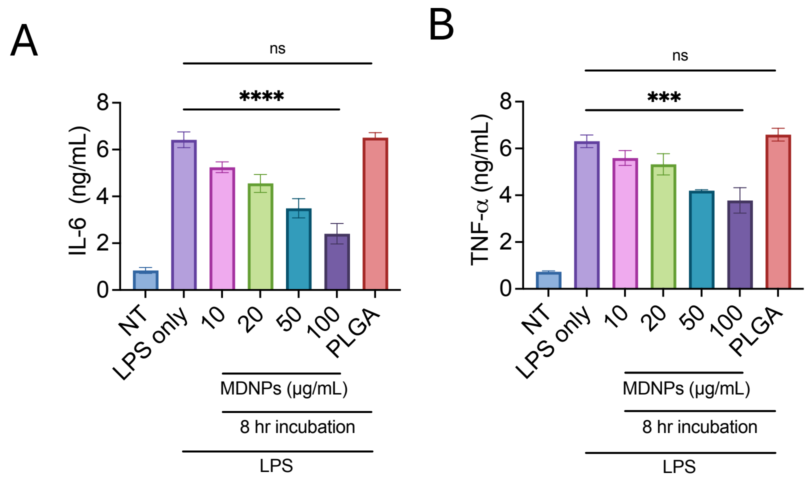


**Supplementary Figure S6.** Concentration-dependent therapeutic test demonstrated that 100 ug/mL exhibited the most significant reduction at 8 hr time point of (A) IL-6 and (B) TNF-α.

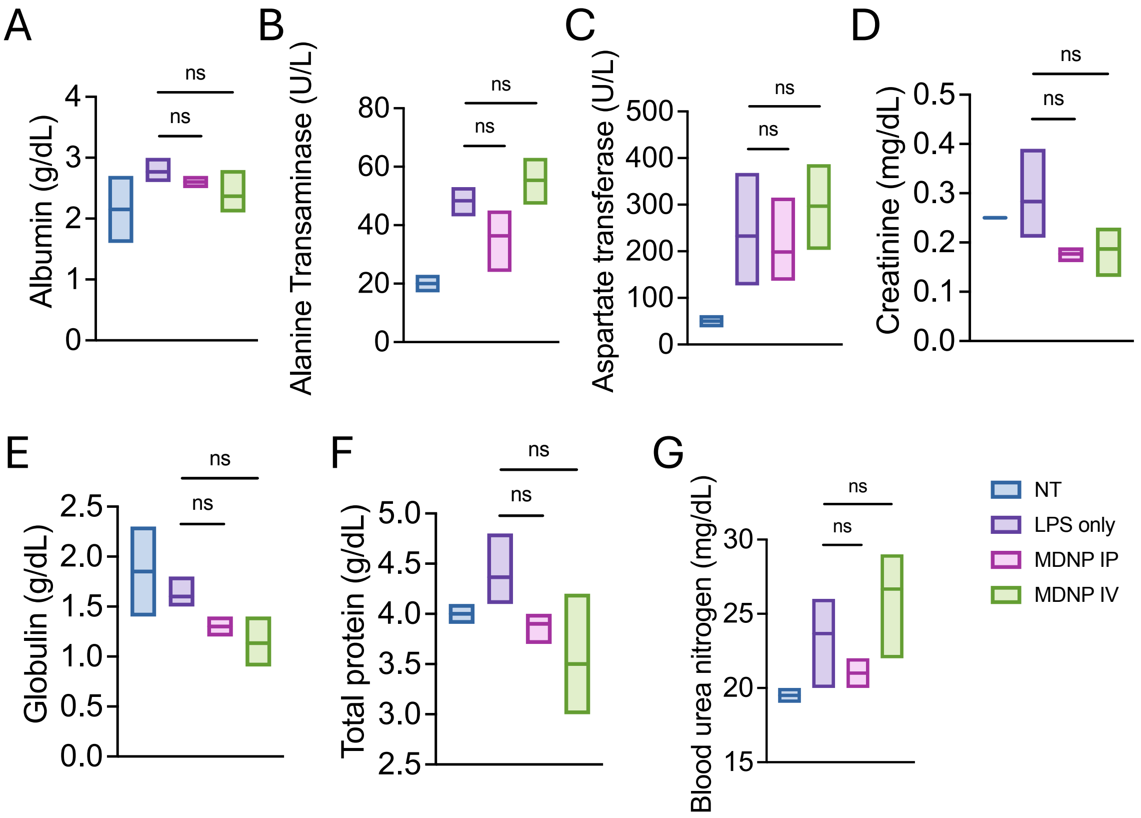


**Supplementary Figure S7**. Blood biochemistry analysis of LPS challenged and MDNPs administered mice plasma (A) Albumin (B) Alanine Transaminase (C) Creatine (D) Aspartate transferase (E) Globulin (F) Total protein (G) Blood Urea Nitrogen. All data is expressed as mean ± SD (n=3/group).

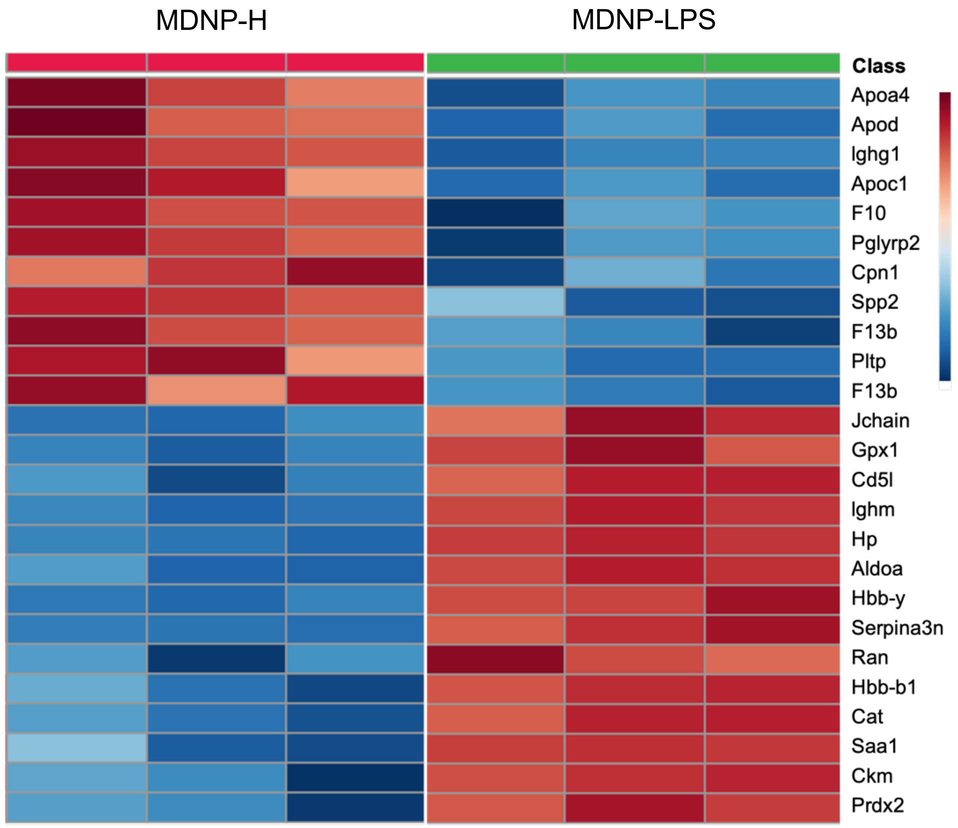


**Supplementary Figure S8.** Top 25 proteins that are upregulated and downregulated in each healthy and LPS plasma incubated group.

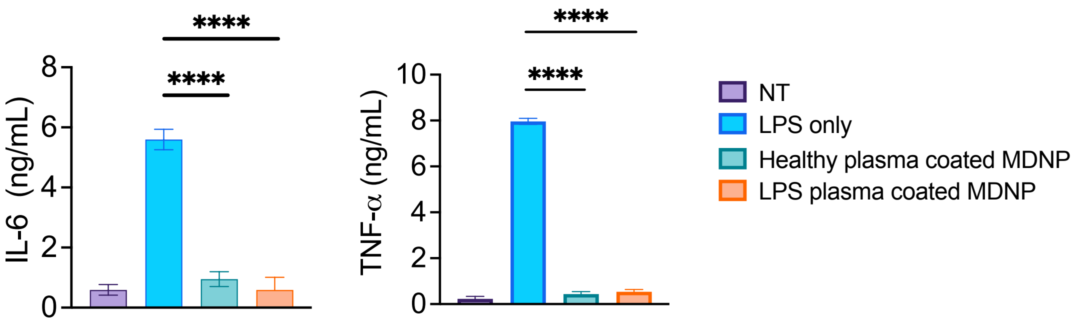


**Supplementary Figure S9.** LPS-induced mouse plasma coated MDNP was tested to show there was no induction of pro-inflammatory cytokines production in BMDM. All data are expressed as means ± SD (n=3). ****$p<0.0005$ versus LPS only group.

| Accession | Description | Gene Symbol | Abundances (Normalized) MDNP-H | Abundances (Normalized) MDNP-LPS |
|---|---|---|---|---|
| Q61171 | Peroxiredoxin-2 | Prdx2 | 3085888.231 | 12099945.33 |
| P03953 | Complement factor D | Cfd | 12019059.35 | 11739969.15 |
| P46412 | Glutathione peroxidase 3 | Gpx3 | 18460002.92 | 11098693.44 |
| P02535 | Keratin, type I cytoskeletal 10 | Krt10 | 9677242.7 | 10915950.63 |
| Q64726 | Zinc-alpha-2-glycoprotein | Azgp1 | 10338349.62 | 10709609 |
| P11859 | Angiotensinogen | Agt | 7796111.559 | 10355122.87 |
| Q00724 | Retinol-binding protein 4 | Rbp4 | 5845338.528 | 9707910.377 |
| Q3UV17 | Keratin, type II cytoskeletal 2 oral | Krt76 | 3509402.221 | 9209172.535 |
| P01592 | Immunoglobulin J chain | Jchain | 2866037.289 | 9103479.2 |
| Q02105 | Complement C1q subcomponent subunit C | C1qc | 8399420.657 | 8647158.17 |
| Q9Z1R3 | Apolipoprotein M | Apom | 7701827.69 | 8550116.051 |
| P31532 | Serum amyloid A-4 protein | Saa4 | 7586819.857 | 8415684.275 |
| Q3TTY5 | Keratin, type II cytoskeletal 2 epidermal | Krt2 | 4551184.715 | 7929576.352 |
| P49182 | Heparin cofactor 2 | Serpind1 | 8334838.569 | 7270216.039 |
| P11589 | Major urinary protein 2 | Mup2 | 3056383.124 | 7112354.108 |
| P51910 | Apolipoprotein D | Apod | 7550189.187 | 7075306.394 |
| Q5FW60 | Major urinary protein 20 | Mup20 | 2352747.672 | 6801889.379 |
| Q61704 | Inter-alpha-trypsin inhibitor heavy chain H3 | Itih3 | 6604642.255 | 6787648.943 |
| Q61730 | Interleukin-1 receptor accessory protein | Il1rap | 7482934.893 | 6706985.092 |
| P42703 | Leukemia inhibitory factor receptor | Lifr | 5740631.518 | 6664111.113 |
| P13634 | Carbonic anhydrase 1 | Ca1 | 2848827.054 | 6608731.101 |
| Q9JJN5 | Carboxypeptidase N catalytic chain | Cpn1 | 8100394.557 | 6459556.99 |
| P35441 | Thrombospondin-1 | Thbs1 | 101055.5742 | 6411564.188 |
| O88947 | Coagulation factor X | F10 | 11204744.52 | 6302198.774 |
| P01887 | Beta-2-microglobulin | B2m | 6897094.775 | 6141391.565 |
| P14847 | C-reactive protein | Crp | 3275921.805 | 5828717.652 |
| P16858 | Glyceraldehyde-3-phosphate dehydrogenase | Gapdh | 1017889.26 | 5791969.956 |
| P70274 | Selenoprotein P | Selenop | 5075468.562 | 5714927.268 |
| Q80YC5 | Coagulation factor XII | F12 | 5250962.792 | 5281776.716 |
| P98086 | Complement C1q subcomponent subunit A | C1qa | 5422527.892 | 5262030.37 |
| Q9Z126 | Platelet factor 4 | Pf4 | 104407.6073 | 5059918.916 |
| P16301 | Phosphatidylcholine-sterol acyltransferase | Lcat | 4408647.341 | 4968372.195 |
| P14106 | Complement C1q subcomponent subunit B | C1qb | 4875764.818 | 4958952.405 |
| O88783 | Coagulation factor V | F5 | 2642773.202 | 4874498.276 |
| Q8R121 | Protein Z-dependent protease inhibitor | Serpina10 | 4870612.494 | 4802069.979 |
| P55065 | Phospholipid transfer protein | Pltp | 4973443.774 | 4531456.258 |
| P01727 | Ig lambda-1 chain V region S43 | | 4235723.765 | 4531136.166 |
| P01655 | Ig kappa chain V-III region PC 7132 | | 2173667.333 | 4148828.728 |
| Q923D2 | Flavin reductase (NADPH) | Blvrb | 1210426.717 | 4053489.672 |
| Q61268 | Apolipoprotein C-IV | Apoc4 | 5251213.727 | 4032759.88 |
| P04945 | Ig kappa chain V-VI region NQ2-6.1 | | 645056.2321 | 4026117.531 |
| Q9R098 | Hepatocyte growth factor activator | Hgfac | 5913123.193 | 3968114.69 |
| P41317 | Mannose-binding protein C | Mbl2 | 3274967.556 | 3947541.403 |
| P01631 | Ig kappa chain V-II region 26-10 | | 8099471.584 | 3928048.55 |
| P01801 | Ig heavy chain V-III region J606 | | 1864864.172 | 3764640.681 |
| Q60994 | Adiponectin | Adipoq | 5218288.283 | 3592772.863 |
| P07310 | Creatine kinase M-type | Ckm | 93710.93473 | 3509827.392 |
| P01843 | Ig lambda-1 chain C region | | 3421048.039 | 3504298.725 |
| P01803 | Ig heavy chain V region AMPC1 | | 587200.1683 | 3323377.141 |
| P04247 | Myoglobin | Mb | 111294.5766 | 3188757.356 |
| P11680 | Properdin | Cfp | 4110900.538 | 3140100.888 |
| P04939 | Major urinary protein 3 | Mup3 | 1602452.567 | 3092342.071 |
| P97298 | Pigment epithelium-derived factor | Serpinf1 | 2630407.051 | 3033494.276 |
| P01680 | Ig kappa chain V-IV region S107B | | 827532.5022 | 2963434.332 |
| Q61508 | Extracellular matrix protein 1 | Ecm1 | 2409400.968 | 2826543.008 |
| Q8VED5 | Keratin, type II cytoskeletal 79 | Krt79 | 1212706.798 | 2776684.772 |
| P01635 | Immunoglobulin kappa chain variable 12-41 | Igkv12-41 | 427261.8639 | 2736059.576 |
| Q02357 | Ankyrin-1 | Ank1 | 36773.72166 | 2629400.818 |
| P15508 | Spectrin beta chain, erythrocytic | Sptb | 506477.6171 | 2626512.316 |
| P08032 | Spectrin alpha chain, erythrocytic 1 | Spta1 | 452189.6159 | 2593533.771 |
| P01665 | Ig kappa chain V-III region PC 7043 | | 171570.79 | 2580106.965 |
| P04918 | Serum amyloid A-3 protein | Saa3 | 635270.9355 | 2457250.179 |
| Q8BH61 | Coagulation factor XIII A chain | F13a1 | 6117289.427 | 2293677.485 |
| Q91WP0 | Mannan-binding lectin serine protease 2 | Masp2 | 2541102.696 | 2285064.866 |
| P01878 | Ig alpha chain C region | | 3224291.133 | 2237437.758 |
| P10126 | Elongation factor 1-alpha 1 | Eef1a1 | 441537.694 | 2159080.241 |
| P70389 | Insulin-like growth factor-binding protein complex acid labile subunit | Igfals | 2224727.582 | 2081860.873 |
| P08071 | Lactotransferrin | Ltf | 28203.9923 | 2037189.124 |
| P18528 | Ig heavy chain V region 6.96 | | 1520840.757 | 1955728.595 |
| P62737 | Actin, aortic smooth muscle | Acta2 | 488055.7673 | 1905757.552 |
| P01644 | Ig kappa chain V-V region HP R16.7 | | 2311965.542 | 1845398.545 |
| P62983 | Ubiquitin-ribosomal protein eS31 fusion protein | Rps27a | 654733.5784 | 1728212.099 |
| Q8VCS0 | N-acetylmuramoyl-L-alanine amidase | Pglyrp2 | 1968563.854 | 1631723.507 |
| P09581 | Macrophage colony-stimulating factor 1 receptor | Csf1r | 1299863.116 | 1629887.006 |
| P82198 | Transforming growth factor-beta-induced protein ig-h3 | Tgfbi | 206323.6036 | 1618988.451 |
| P24270 | Catalase | Cat | 237052.1895 | 1575287.396 |
| P01638 | Ig kappa chain V-V region L6 | | 1079440.707 | 1533036.36 |
| P02104 | Hemoglobin subunit epsilon-Y2 | Hbb-y | 355490.4868 | 1498424.351 |
| Q62351 | Transferrin receptor protein 1 | Tfrc | 987638.7793 | 1469745.1 |
| Q9JHH6 | Carboxypeptidase B2 | Cpb2 | 1539544.598 | 1450523.22 |
| P05064 | Fructose-bisphosphate aldolase A | Aldoa | 183641.871 | 1446645.03 |
| Q07968 | Coagulation factor XIII B chain | F13b | 3531441.262 | 1427418.032 |
| P01639 | Immunoglobulin kappa chain variable 9-120 | Igkv9-120 | 1482319.534 | 1391522.029 |
| P15327 | Bisphosphoglycerate mutase | Bpgm | 517656.4183 | 1388413.446 |
| P01786 | Ig heavy chain V region MOPC 47A | | 378980.9374 | 1346201.385 |
| P61939 | Thyroxine-binding globulin | Serpina7 | 1050672.157 | 1306869.921 |
| P01897 | H-2 class I histocompatibility antigen, L-D alpha chain | H2-L | 782551.5554 | 1305391.625 |
| P39039 | Mannose-binding protein A | Mbl1 | 804163.9994 | 1292026.996 |
| O70165 | Ficolin-1 | Fcn1 | 989373.7446 | 1289473.009 |
| P33587 | Vitamin K-dependent protein C | Proc | 918418.4606 | 1245425.217 |

| Accession | Description | Gene Symbol | Abundances (Normalized) MDNP-H | Abundances (Normalized) MDNP-LPS |
|---|---|---|---|---|
| P01844 | Ig lambda-2 chain C region | Iglc2 | 933514.4961 | 1199107.648 |
| P39876 | Metalloproteinase inhibitor 3 | Timp3 | | 1198696.864 |
| P11352 | Glutathione peroxidase 1 | Gpx1 | 448819.0804 | 1156240.245 |
| Q02013 | Aquaporin-1 | Aqp1 | 290921.9173 | 1152957.361 |
| P01787 | Ig heavy chain V regions | | 773081.5033 | 1152956.555 |
| P51437 | Cathelicidin antimicrobial peptide | Camp | 71690.20351 | 1141140.866 |
| P18525 | Ig heavy chain V region 5-84 | | 929417.0169 | 1113167.285 |
| Q61805 | Lipopolysaccharide-binding protein | Lbp | 212012.418 | 1106587.479 |
| P06151 | L-lactate dehydrogenase A chain | Ldha | 83259.45611 | 1068558.922 |
| Q8BPB5 | EGF-containing fibulin-like extracellular matrix protein 1 | Efemp1 | 957110.2576 | 1051273.401 |
| P18524 | Ig heavy chain V region RF | | 996843.9067 | 1027726.632 |
| Q9CQW3 | Vitamin K-dependent protein Z | Proz | 1475605.916 | 1018596.468 |
| Q8CG16 | Complement C1r-A subcomponent | C1ra | 1072269.264 | 1007713.106 |
| P01758 | Ig heavy chain V region 108A | Igh-VJ558 | 311409.0389 | 1003340.943 |
| O35930 | Platelet glycoprotein Ib alpha chain | Gp1ba | 1210609.365 | 973056.378 |
| P17742 | Peptidyl-prolyl cis-trans isomerase A | Ppia | 114314.0033 | 962190.9382 |
| P21180 | Complement C2 | C2 | 974672.7363 | 915365.3758 |
| P52480 | Pyruvate kinase PKM | Pkm | 66810.24807 | 768374.2116 |
| P02089 | Hemoglobin subunit beta-2 | Hbb-b2 | | 762368.0511 |
| P01643 | Ig kappa chain V-V region MOPC 173 | | 676756.4934 | 759604.6229 |
| P11404 | Fatty acid-binding protein, heart | Fabp3 | 49282.86044 | 751989.3473 |
| Q9WVJ3 | Carboxypeptidase Q | Cpq | 481426.3933 | 745130.3249 |
| P63017 | Heat shock cognate 71 kDa protein | Hspa8 | 452344.8245 | 725191.1799 |
| Q6IMP4 | Pannexin-2 | Panx2 | 673329.6096 | 714597.5646 |
| P62827 | GTP-binding nuclear protein Ran | Ran | 240544.5144 | 709241.6649 |
| P17751 | Triosephosphate isomerase | Tpi1 | 59028.4323 | 681226.2498 |
| P62806 | Histone H4 | H4 | | 650320.7572 |
| P08228 | Superoxide dismutase [Cu-Zn] | Sod1 | 284304.0683 | 633104.1819 |
| P70663 | SPARC-like protein 1 | Sparcl1 | 59116.26748 | 620773.748 |
| P01662 | Ig kappa chain V-III region ABPC 22/PC 9245 | | 243595.2228 | 598474.2129 |
| P00687 | Alpha-amylase 1 | Amy1 | 636575.4654 | 593350.912 |
| P98064 | Mannan-binding lectin serine protease 1 | Masp1 | 700603.2732 | 563752.9026 |
| P01750 | Ig heavy chain V region 102 | | 714952.629 | 551904.1395 |
| P10810 | Monocyte differentiation antigen CD14 | Cd14 | | 545869.5645 |
| P11672 | Neutrophil gelatinase-associated lipocalin | Lcn2 | | 535419.015 |
| P14152 | Malate dehydrogenase, cytoplasmic | Mdh1 | | 534558.042 |
| O88968 | Transcobalamin-2 | Tcn2 | 474797.4922 | 529356.3326 |
| P10605 | Cathepsin B | Ctsb | 369116.9063 | 507950.2788 |
| O08899 | Regulator of G-protein signaling 4 | Rgs4 | 508507.6258 | 499218.1259 |
| P01642 | Ig kappa chain V-V region L7 | Gm10881 | 495826.2892 | 496242.8161 |
| P12246 | Serum amyloid P-component | Apcs | | 489876.1313 |
| Q9ET66 | Peptidase inhibitor 16 | Pi16 | 95961.47752 | 479325.348 |
| P35700 | Peroxiredoxin-1 | Prdx1 | 117636.8253 | 478144.7418 |
| P14430 | H-2 class I histocompatibility antigen, Q8 alpha chain | H2-Q8 | 239335.4636 | 469309.8116 |
| Q08879 | Fibulin-1 | Fbln1 | 419877.2255 | 426677.1854 |
| P53986 | Monocarboxylate transporter 1 | Slc16a1 | 28623.60822 | 412682.9054 |
| P27661 | Histone H2AX | H2ax | 109172.7495 | 382814.2872 |
| Q8K1I3 | Secreted phosphoprotein 24 | Spp2 | 338014.7574 | 374289.839 |
| P23492 | Purine nucleoside phosphorylase | Pnp | 254767.5027 | 365869.5486 |
| P05208 | Chymotrypsin-like elastase family member 2A | Cela2a | 80332.75844 | 363113.5375 |
| P48193 | Protein 4.1 | Epb41 | 103875.1831 | 344479.818 |
| P06327 | Ig heavy chain V region VH558 A1/A4 | Gm5629 | 108375.0834 | 332725.2552 |
| P01633 | Immunoglobulin kappa chain variable 6-17 | Igkv6-17 | 197004.8172 | 322070.4402 |
| P01728 | Ig lambda-2 chain V region | | 217489.6166 | 319542.4043 |
| P04117 | Fatty acid-binding protein, adipocyte | Fabp4 | | 315778.6717 |
| P16015 | Carbonic anhydrase 3 | Ca3 | 22174.50082 | 290487.8667 |
| Q61330 | Contactin-2 | Cntn2 | 134956.1178 | 289166.3145 |
| O55042 | Alpha-synuclein | Snca | 168409.0136 | 282600.449 |
| P10639 | Thioredoxin | Txn | 114151.1231 | 282118.4762 |
| P32848 | Parvalbumin alpha | Pvalb | | 281275.3964 |
| P70296 | Phosphatidylethanolamine-binding protein 1 | Pebp1 | | 239032.0395 |
| P01630 | Ig kappa chain V-II region 7S34.1 | | 280837.7285 | 220323.3254 |
| Q3SXB8 | Collectin-11 | Colec11 | 167401.0377 | 220086.0044 |
| P01902 | H-2 class I histocompatibility antigen, K-D alpha chain | H2-K1 | 96439.86213 | 208128.1475 |
| O55028 | Branched-chain alpha-ketoacid dehydrogenase kinase | Bckdk | 736090.4191 | 206778.6285 |
| Q62009 | Periostin | Postn | 152431.3956 | 201674.5953 |
| Q8K426 | Resistin-like gamma | Retnlg | 60400.40284 | 197528.973 |
| P29533 | Vascular cell adhesion protein 1 | Vcam1 | 154212.4878 | 195456.0182 |
| Q8R0Z6 | Angiopoietin-related protein 6 | Angptl6 | 148958.7772 | 194914.202 |
| Q9QUM9 | Proteasome subunit alpha type-6 | Psma6 | 102363.4692 | 193835.0291 |
| P43025 | Tetranectin | Clec3b | 394119.509 | 192169.469 |
| Q9D358 | Low molecular weight phosphotyrosine protein phosphatase | Acp1 | 74136.96662 | 186169.208 |
| Q99PT1 | Rho GDP-dissociation inhibitor 1 | Arhgdia | | 185554.6829 |
| P16294 | Coagulation factor IX | F9 | 205489.3198 | 176598.7952 |
| O08692 | Neutrophilic granule protein | Ngp | | 173644.8302 |
| P01660 | Ig kappa chain V-III region PC 3741/TEPC 111 | | 451258.3076 | 172938.9516 |
| P26928 | Hepatocyte growth factor-like protein | Mst1 | 220308.4434 | 165340.6285 |
| P62259 | 14-3-3 protein epsilon | Ywhae | 25740.68703 | 165025.5098 |
| Q64339 | Ubiquitin-like protein ISG15 | Isg15 | | 155448.3283 |
| P01819 | Ig heavy chain V region MOPC 141 | | 111665.7851 | 153986.9175 |

| Accession | Description | Gene Symbol | Abundances (Normalized) MDNP-H | Abundances (Normalized) MDNP-LPS |
|---|---|---|---|---|
| P13597 | Intercellular adhesion molecule 1 | Icam1 | 35830.14385 | 153049.0449 |
| Q8BK48 | Pyrethroid hydrolase Ces2e | Ces2e | 138648.6251 | 141534.1001 |
| Q9DA19 | Corepressor interacting with RBPJ 1 | Cir1 | 143355.989 | 131390.7276 |
| Q03311 | Cholinesterase | Bche | 17536.47352 | 129654.3962 |
| P09411 | Phosphoglycerate kinase 1 | Pgk1 | | 128763.9831 |
| P48774 | Glutathione S-transferase Mu 5 | Gstm5 | | 128138.7795 |
| P45700 | Mannosyl-oligosaccharide 1,2-alpha-mannosidase IA | Man1a1 | 111866.5159 | 125361.5957 |
| P70290 | 55 kDa erythrocyte membrane protein | Mpp1 | | 120048.7397 |
| O08709 | Peroxiredoxin-6 | Prdx6 | 45497.51336 | 117590.9447 |
| O70435 | Proteasome subunit alpha type-3 | Psma3 | 56781.75446 | 111797.9731 |
| P01674 | Ig kappa chain V-III region PC 2154 | | 180993.5702 | 106219.4872 |
| P07901 | Heat shock protein HSP 90-alpha | Hsp90aa1 | | 100481.8017 |
| P54116 | Stomatin | Stom | | 98450.13141 |
| P21460 | Cystatin-C | Cst3 | 53373.02358 | 94348.54252 |
| P01636 | Ig kappa chain V-V region MOPC 149 | | 68964.84947 | 92414.91495 |
| P62962 | Profilin-1 | Pfn1 | 25524.30285 | 91376.00012 |
| P01741 | Ig heavy chain V region | | 165123.7841 | 88549.54136 |
| P10853 | Histone H2B type 1-F/J/L | | | 85259.87042 |
| P30412 | Peptidyl-prolyl cis-trans isomerase C | Ppic | 54402.95832 | 84604.76957 |
| Q8CIF4 | Biotinidase | Btd | 72066.23784 | 82305.05012 |
| O89103 | Complement component C1q receptor | Cd93 | | 75276.41839 |
| Q05020 | Apolipoprotein C-II | Apoc2 | 97660.29413 | 67462.46407 |
| Q60805 | Tyrosine-protein kinase Mer | Mertk | 61380.59791 | 67195.69242 |
| P18337 | L-selectin | Sell | 58937.10843 | 66985.2593 |
| Q9R1P4 | Proteasome subunit alpha type-1 | Psma1 | 50646.0636 | 66984.96607 |
| Q8CFG8 | Complement C1s-1 subcomponent | C1s2 | 62510.4846 | 55936.92934 |
| Q01853 | Transitional endoplasmic reticulum ATPase | Vcp | 15995.53715 | 48063.65088 |
| P28666 | Murinoglobulin-2 | Mug2 | | 48007.14176 |
| P49222 | Protein 4.2 | Epb42 | | 38722.11108 |
| P09470 | Angiotensin-converting enzyme | Ace | 54092.72918 | 36537.61521 |
| Q8BFZ3 | Beta-actin-like protein 2 | Actbl2 | | 35956.44498 |
| Q9DCD0 | 6-phosphogluconate dehydrogenase, decarboxylating | Pgd | 14439.63011 | 33679.13632 |
| Q8BU03 | Periodic tryptophan protein 2 homolog | Pwp2 | 793572.6938 | 28688.57962 |
| Q8VDD5 | Myosin-9 | Myh9 | | 28645.33948 |
| P63242 | Eukaryotic translation initiation factor 5A-1 | Eif5a | | 18835.75916 |
| Q01755 | T-complex protein 11 | Tcp11 | 16377.45733 | 18424.59229 |
| Q14B46 | Rhotekin-2 | Rtkn2 | 2911346.75 | |

**Supplementary Table S1.** A complete list of total 297 protein corona in healthy and LPS group with abundance.
